# A treasure trove of 1,034 actinomycete genomes

**DOI:** 10.1101/2024.01.16.574955

**Authors:** Tue Sparholt Jørgensen, Omkar Mohite, Eva B Sterndorff, Maria Alvarez-Arevalo, Kai Blin, Thomas J Booth, Pep Charusanti, David Faurdal, Troels Ø Hansen, Matin Nuhamunada, Anna-Sophie Mourched, Bernhard Ø Palsson, Tilmann Weber

## Abstract

Filamentous Actinobacteria, recently renamed Actinomycetia, are the most prolific source of microbial bioactive natural products. Studies on biosynthetic gene clusters benefit from or require chromosome-level assemblies. Here, we provide DNA sequences from more than 1,000 isolates: 881 complete genomes and 153 near-complete genomes, representing 28 genera and 389 species, including 244 likely novel species. All genomes are from filamentous isolates of the class Actinomycetia from the NBC culture collection. The largest genus is *Streptomyces* with 886 genomes including 742 complete assemblies. We use this data to show that analysis of complete genomes can bring biological understanding not previously derived from more fragmented sequences or less systematic datasets. We document the central and structured location of core genes and distal location of specialized metabolite biosynthetic gene clusters and duplicate core genes on the linear *Streptomyces* chromosome, and analyze the content and length of the terminal inverted repeats which are characteristic for *Streptomyces*. We then analyze the diversity of trans-AT polyketide synthase biosynthetic gene clusters, which encodes the machinery of a biotechnologically highly interesting compound class. These insights have both ecological and biotechnological implications in understanding the importance of high quality genomic resources and the complex role synteny plays in Actinomycetia biology.

## Introduction

The bacterial class Actinomycetia (recently reclassified as Actinobacteria) is hugely successful, with species commonly inhabiting a wide range of niches, both host-associated and environmental. Environmental Actinomycetia species, especially those inhabiting soil, often have a complex biology and a filamentous growth pattern, which makes laboratory studies difficult.

In the past century, there has been great scientific interest surrounding the class Actinomycetia. Many essential medicines such as antibiotics (e.g., tetracycline, erythromycin and vancomycin), immunosuppressants (e.g., rapamycin), anthelmintics (e.g., avermectin), and anticancer drugs (e.g. actinomycin D and daunorubicin), but also important insecticides (e.g., spinosyns) and food preservatives (e.g., ε-polylysine) either directly come from or are derivatives of compounds produced by Actinomycetia strains ^1, 2^.

The biosynthesis of these natural products, often referred to as secondary or specialized metabolites, is encoded in genomic regions called biosynthetic gene clusters (BGCs). These BGCs code for a variety of enzymes that catalyze the biosynthesis of highly complex chemicals from “simple” precursors from primary metabolism. In addition to the biosynthetic function, the BGCs often also contain genes for regulation^3^, molecular transport/export^4^, and self-resistance^5^. Due to a high degree of conservation of the key enzymatic steps in the biosynthesis of these specialized metabolites, regions of the genome encoding such BGCs can be identified and annotated using bioinformatics tools such as antiSMASH^6^, which classifies a BGC based on a set of rules specific to each BGC type^7^. AntiSMASH then compares the identified BGC to both known specialized metabolite pathways and existing genomic sequences, and suggests possible precursors. These features have established genome mining as a complementary approach to the bioactivity and compound-centered discovery of novel bioactive molecules^8^. As some of these BGCs, such as the gargantulide BGC^9^, have sizes of more than 200 kbp, high quality genome assemblies have a strong influence on the quality of the genome mining analyses, as contiguous sequence data allows the identification of complete gene clusters and not just fragments – a challenge that is prevalent with low to medium quality draft genome and many metagenomic datasets^10^.

Most bacterial genomes consist of a single circular chromosome and maybe some smaller circular plasmids. Some genera of the Actinomycetia also have circular chromosomes, but members of the genus *Streptomyces* have large, linear genomes, similar to how most eukaryotic genomes are organized. The linear *Streptomyces* genome is usually capped with terminal inverted repeats (TIRs)^11^, which can be more than 1 Mbp long and can encode BGCs (e.g. ^12^). The TIRs contain short telomeres which are capped with covalently bound terminal proteins^13^. The TIRs are highly dynamic and have been speculated to play a role in gene dose modulation by duplicating genomic regions, thereby potentially increasing compound production or diversity if BGCs are duplicated^14^. The TIRs also present a special challenge in genome assembly, as even long read information regularly is not of sufficient length to cover the TIR from the chromosome start to the most terminal non-repeat regions. However, because of the repetitive nature, it is still possible to resolve TIRs and generate complete genome sequences. Information about telomeres, TIRs and chromosome arms is not a commonly recorded statistic for bacterial genome assemblies. It is possible that the duplicate TIRs are missing, placed on their own contig, or otherwise misrepresented in otherwise complete genome sequences of *Streptomyces* isolates in public databases, as large scale errors relating to the TIRs are often observed artifacts introduced by sequence assembly software optimized for circular bacterial genomes. The linear *Streptomyces* genome is widely hypothesized to be composed of a central ‘core’, where the basic cellular functions are encoded, and distal ‘arms,’ which are dynamic and more likely to encode BGCs and accessory functions and often contain mobile genetic elements. Several studies have reported this genomic compartmentalization in smaller datasets^15^ or using e.g. rRNA genes to mark the core region^16^.

Previously, the high GC content (65-75 %) of many Actinomycetia species hampered the ability to sequence complete genomes. But with the advent of the GC-agnostic nanopore sequencing technology^17^, and implementation of lab protocols minimizing GC bias from illumina sequences^18^, high GC-content is less of an obstacle. It is now possible to obtain complete genomes using technologies such as PacBio or Nanopore DNA sequencing, but currently available genomic resources are still too scarce to cover the biological diversity of Actinomycetia in nature^19^, and large-scale efforts are often focused on quantity over completeness and chromosomal organization of the individual genome. In a recent massive collaborative study involving both the U.S. Department of Energy Joint Genome Institute (JGI) and the German Collection of Microorganisms and Cell Cultures (DMSZ), the lack of high-quality resources was suggested to severely limit both the biological understanding of Actinomycetia species and the development of new bioactive compounds from this clade^19^.

We report the whole genome sequencing of 1,034 Actinomycetia strains from the New Bioactive Compounds (NBC) Collection of the Novo Nordisk Foundation Center for Biosustainability at the Technical University of Denmark; 881 strains with complete genomes and 153 near-complete genomes. We provide analysis of their phylogeny, genomic organization, BGC content and provide an example of how this large dataset can be used to identify novel BGCs.

## Materials & Methods

For basic statistics on the G1034 genomes, please refer to Supplementary Material S1

### Biological Resources: Isolation, growth and DNA extraction of the NBC collection

Soil samples were collected by digging approximately 7-10 cm below the surface and placing a small scoop of soil into plastic bags or 50 mL Falcon tubes. Once back in the lab, the samples were stored at 4 °C for a maximum of 3 weeks before processing. Detailed metadata of the samples can be found in the individual [BioSample](https://www.ncbi.nlm.nih.gov/bioproject/?term=PRJNA747871) records.

To isolate actinomycete strains, we mixed 1 g of soil with 3 mL 6 % yeast extract, incubated the sample in a 60 °C water bath for 20 min, and then diluted this mixture 1:49 with sterile water. We then spread 200 μL of this dilution onto 3 types of agar media: ISP4, AIM-Avicel, and AIM-xylose in 94 x 16 mm petri dishes containing approximately 25 mL of agar media. No antifungal compounds or inhibitors against Gram-negative bacteria were used. The plates were incubated at 30 °C. After 5, 10, 15 and 20 days incubation, we picked single colonies and restreaked them onto ISP2 plates. These plates were again incubated at 30 °C. Once actinomycete growth was observed on the ISP2 plates, a chunk of the bacteria was lifted from the agar and placed into triple-baffled flasks containing ISP2 liquid media. The cultures were incubated at 30 °C with shaking. After robust growth, an aliquot was mixed 1:1 with 50 % glycerol to make frozen stocks, which were stored at −80 °C.

A detailed description of the growth conditions and DNA extraction protocol can be found in ^18^. Briefly, strains were grown in liquid ISP2 or YEME medium in baffled flasks at 30 °C with constant stirring. Then the cells were harvested and washed with PBS, and DNA extracted with a modified Qiagen Genomic Tip 20 protocol. For strains with an aggregated growth pattern, physical shearing in liquid nitrogen using a mortar and pestle was applied. The concentration, purity, and amount of short DNA fragments of the resulting purified DNA was evaluated with Qubit BR dsDNA, nanodrop, and 0.7 % agarose gel electrophoresis. The length of DNA fragments using this method is generally around 50,000 bp^18^. Samples with many short DNA fragments were further purified using either the Short Read Eliminator kit (PacBio, San Diego, CA, USA), or 0.4 - 0.5 x volume KAPA PURE SPRI beads (Roche, Basel, Switzerland).

### Illumina sequencing

Illumina data was either generated in-house on an Illumina NextSeq500 with the KAPA HyperPlus protocol (Roche, Basel, Switzerland), or at Novogene Inc (Beijing, China) on an Illumina Novaseq machine with the libraries either prepared using the NEBNext® Ultra™ II DNA Library Prep Kit or the Novogene NGS DNA Prep Set kit, both latter using a special 6 PCR cycle protocol to minimize GC bias in the sequencing results. All libraries were sequenced using 2×150 nt paired end sequencing kits. Raw illumina reads were trimmed with Trim Galore v0.6.4_dev ^20^ in paired mode with the minimum length set to 100 bp and the quality cutoff set to Q20.

### Nanopore sequencing

Nanopore sequencing was performed on the Nanopore MinION platform, with either the SQK-RBK004 or SQK-RBK110-96 kit, both of which are transposase based. The protocol was modified as described in ^18^. Briefly, the DNA to transposase ratio was modified to allow for longer fragments and a post tagmentation size selection was performed to remove small fragments. All basecalling was performed with Guppy (v.5.0.17+99baa5b, client-server API version 7.0.0), which also was used for demultiplexing and adapter trimming.

### Assembly and preliminary analysis

The workflow AAA-Actino-Assembly-and-Annotation (v.1.6.7) was used for all assembly and preliminary analysis (https://github.com/tuspjo/AAA-Actino-Assembly-and-Annotation). Default parameters were used when not otherwise mentioned. The workflow consists of the following steps:

The nanopore reads were assembled using Flye (v.2.9)^21^ with the following settings: 5 polishing iterations and using the --nano-raw switch. The N50 and total number of bases in reads were extracted from the Flye statistics after assembly, and thus represent the dataset used for the assembly. The assembly graphs were manually inspected in Bandage v.0.9.0^22^, and in cases where Flye was unable to completely resolve an unambiguous graph, we applied the simplistic NPGM-contigger (https://github.com/biosustain/npgm-contigger). The topology of each chromosome is reported based on the manual inspection of the repeat graph and of the contigging. In cases where the assembly graph was ambiguous, we first simplified the dataset using Filtlong (https://github.com/rrwick/Filtlong) to remove the shortest and lowest quality reads according to Supplementary Material S1, and if a complete genome was still not obtained, deposited the genome as the lower contiguity “WGS” rather than “chromosome”. Coverage plots of both illumina (Bowtie2^23^) and nanopore (minimap2^24^) data mappings were manually inspected for sudden coverage changes to half or double the surrounding area coverage (see Supplementary Material S2 for coverage plot examples). The coverage plot inspection revealed large scale assembly artifacts of some of the chromosomes, resulting in 11 genomes which needed to be reassembled after Filtlong removal of half the raw nanopore data. These errors related to the TIRs of *Streptomyces* genomes.

### Sequence sanity checks

Illumina reads were mapped on the nanopore assembly using Bowtie2 v.2.3.4.3 ^23^. This sanity check was failed if less than 60 % of illumina reads mapped, and the sample was excluded from further analysis. This step is taken to ensure that the nanopore and illumina datasets come from the same sample.

### Assembly polishing

All genomes were first polished using the nanopore data with the Flye polisher, this includes both samples contigged by Flye and by the NPGM-contigger. A total of five iterations of nanopore read polishing was used. Then, samples were polished with the illumina reads using Polypolish v.0.5.0^25^, and then using POLCA (part of MaSuRCA v.4.0.5 ^26^), as described in Wick et al.^25^.

### Gene and functional annotation, phylogenetic placement, assembly quality, and antiSMASH annotation

Gene and functional annotation was performed at NCBI using the PGAP pipeline. Phylogenetic assignment was done using GTDB-Tk v.1.0.2 (reference data version r207) ^27^, and later verified at NCBI. In the handful of cases where the taxonomic assignment between NCBI and GTDB did not agree, or a genus was differently named, the NCBI assignment was kept in the genbank annotation. For the strain NBC_01635, GTDB-Tk repeatedly failed. However, based on the presence of core genes, the linear chromosome organization with TIRs, and the NCBI classification, this strain was classified as *Streptomyces sp*.

We used the BUSCO ^28^ program to represent different levels of evolutionarily conserved genes. BUSCO ^28^ uses sets of HMMs of universal (defined as >90 % of species) single-copy orthologous genes to estimate the completeness and duplication levels of eu- and prokaryotic genome assemblies. It has become nearly universally adapted for assessing assemblies, and 193 BUSCO datasets currently exist for domain (e.g Bacteria), phylum (e.g. Actinomycetota), class (e.g. Actinomycetia), order (e.g. Streptomycetales), and more classification levels. BUSCO v.5.1.2 ^28^ with the databases bacteria_odb10 and actinobacteria_class_odb10 was used to estimate the gene placement on the genomes and core gene content, respectively.

AntiSMASH v.7.0.0 ^6^ was used to predict BGCs with the NCBI PGAP annotation in genbank format as input and the following parameters: --cb-general --cb-subclusters --cb-knownclusters --genefinding-tool none --clusterhmmer --cc-mibig --asf --tigr --pfam2go.

### Developed tools and additional used bioinformatics methods

#### *DnaA* flipping

For the analysis of complete *Streptomyces* genomes, all sequences were oriented so that the *dnaA* gene was in the forward direction using the biopython script deposited to https://github.com/kblin/dna-flipper. In total, 368 genomes (49.6 %) were flipped, while 347 (50.4 %) were not.

#### Plasmid identification

We manually curated a list of 170 plasmid specific PFAM domains (deposited to https://github.com/tuspjo/G1000_manuscript_analyses) and used it to classify extrachromosomal elements as plasmids if any of the plasmid-specific PFAM domains were present. This methodology is similar to a previously developed method and classified 85 % of extrachromosomal elements as plasmids ^29^. The accuracy of this methodology was not experimentally confirmed. Elements without plasmid-like traits were classified as “extrachromosomal”.

#### gbk to faa and gff3

For conversion from genbank format file to GFF3, we used bioconvert^30^. For conversion from gbk to amino acid fasta, we used the script at https://www.biostars.org/p/151891/ written by Istvan Albert.

#### Extracting protoclusters from antiSMASH results

The python script used to extract protoclusters from antiSMASH-annotated genbank-files was deposited to https://github.com/dalofa/gbk_protocluster_parse.

#### TIR identification

To investigate the size of linear inverted repeats on the 742 *Streptomyces* linear chromosomes, we performed a BLAST search ^31^ of the *Streptomyces* chromosome level chromosomes against itself with the following settings: blastn v2.10.0+, gapopen 2 gapextend 1, hit starting on the first 99 bp of the query and ending on the last 99 bp of the subject to account for end polishing artifacts.

#### Submission to NCBI

All raw reads used for assembly, as well as the assemblies, were deposited at NCBI under project number PRJNA747871.

#### Basic statistics

For calculating means, medians, quartiles, standard deviations and other basic metrics, GNU datamash was used ^32^. For calculating the GC %, quast (v.5.2.0) was used ^33^.

#### Data visualization: Kernel density plots

To represent each gene and each BGC equally regardless of the length, we used the midpoint position of each gene model or BGC protocluster. This ensures that long and short BGCs have equal weight in the analysis. The Seaborn kernel density estimate plot “kdeplot” was used on core genes, duplicate core genes, and protocluster antiSMASH categories and types, respectively, with a smoothing of 0.5 (bw_adjust=.5). This means that the density of each plot is independent, and as such the number of genes or clusters necessary for a certain density is not constant between plots, so they can not be directly compared, however the position (x-axis) is constant and directly comparable between plots. The smoothing value used was chosen as it was found to minimize noise while maintaining clarity, however especially in the distal parts of the chromosome, artifacts are observed as the density will have a tendency to approach 0 even when the data does not show basis of this. As the 742 complete *streptomyces* genomes each have 124 bacterial BUSCO gene models, and as the secondary BUSCO copy has been filtered out in their own plot, there are 742 observations for each BUSCO gene model, meaning that the area in the density plot is equal for each gene model, except for two genomes which each had 1 BUSCO gene missing, which means that only 741 observations are available for those two BUSCO gene models. For the duplicate core genes kernel density plot and the antiSMASH category plots, the number of observations of each type vary, and the area of each density is proportional to the number of observations, which can then be compared to each other. However, it is important to repeat that the densities can not be compared between subplots in the kernel density plots.

### Comparison of genome quality with respect to the existing public dataset

The genus *Streptomyces* was the most abundant in the presented dataset and thus was selected for comparison of genome quality with respect to existing datasets in the public repositories. We retrieved 2,938 genomes of the Streptomycetaceae family that were present in the NCBI RefSeq database as of 30 June 2023. These genomes were assigned taxonomic definitions according to the GTDB database ^27, 34^. A total of 462 genomes were not assigned to any of the GTDB-defined species. We calculated the MASH^35^ distance metric across these genomes to assess the diversity based on whole genome sequence similarity. We used a cutoff of 95% similarity (0.05 MASH-distance) to reconstruct a genome similarity network. To calculate the diversity we used the Louvain community detection algorithm that assigns different communities in the network. These communities are designated as different MASH-based species (Supplementary Material S1). The genomes belonging to the genus *Streptomyces* were selected for further analysis (Figure 1C).

**Figure 1.**
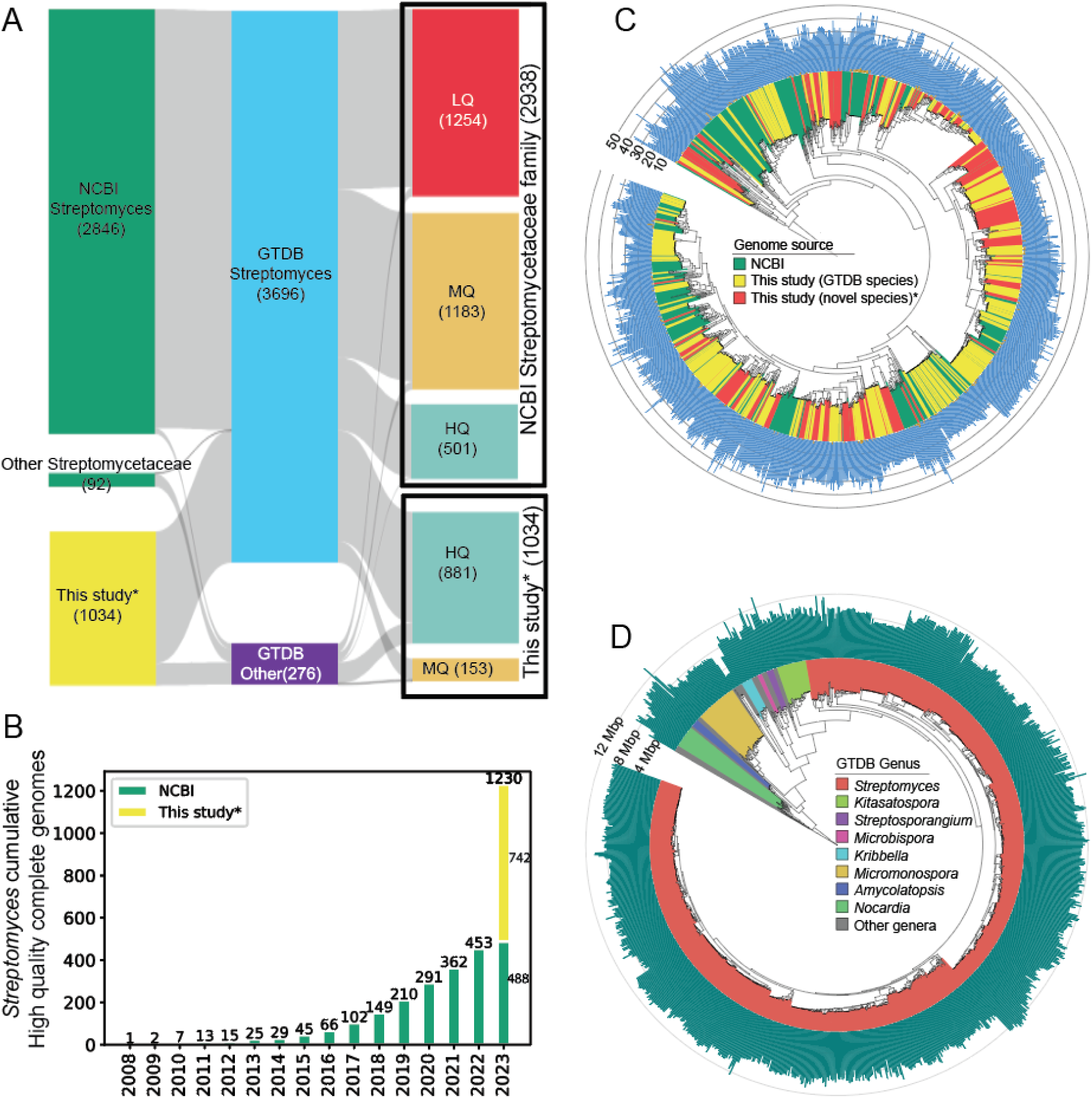
Metadata comparison of *Streptomyces* genomes from NBC collection against publically available genome. A) Taxonomy and quality assessment of 2,938 *Streptomycetaceae* genomes and the 1,034 actinomycetes genomes from this study. The GTDB assignment showed 3,696 of these genomes belonging to the genus *Streptomyces* in total (middle panel of Sankey diagram). B) Cumulative number of HQ *Streptomyces* genomes in the NCBI RefSeq database (as of 30 June, 2023). The yellow bar represents the complete or chromosome level assemblies of *Streptomyces* genomes collected in the present study. The slight difference between the *Streptomyces* numbers in panel A (501) and B (488) are due to differences in the taxonomic assignment between NCBI and GTDB. C) Phylogenetic tree of all high quality complete Streptomyces genomes reconstructed using GTDBTk de novo method. Colors represent the genomes from either NCBI or this study (with or without GTDB species assignment). The bar chart on the outer ring denote the number of BGCs. D) Phylogenetic tree of the 1034 genomes presented in this study, reconstructed using GTDBTk de novo method. Colors represent select genera, whereas the bar chart on the outer ring denotes genome length.

The genomes were classified into three different categories based on the completeness of the assemblies. The complete or chromosome level assemblies were classified as high-quality (HQ), contig or scaffold level assemblies with less than or equal to 100 contigs were classified as medium-quality (MQ), whereas rest were low-quality (LQ). We selected the genomes of high-quality to evaluate the progress of sequencing high-quality genomes of the genus *Streptomyces* over the years and distribution across country of origin (Figures 1A and 1B, see Supplementary material S3 for a list of countries of origin).

### Phylogenetic tree reconstruction

Two different phylogenetic trees were reconstructed here using GTDB-TK ^27^. The dataset of complete genomes of *Streptomyces* contained 1230 genomes with 488 genomes from NCBI and 742 from this study. The NBC_01635 genome was assigned to the genus *Streptomyces* according to NCBI but not GTDB and thus was ignored. The resulting 1229 genomes were processed using GTDB-TK’s de novo tree construction algorithm. *Kitasatospora* GCF_000269985.1 was used as an outgroup. Another tree was reconstructed using all 1,034 genomes presented in this study. For this, *Mycobacterium tuberculosis* GCF_000195955.2 was used as an outgroup. Tree visualizations were generated using iTOL. For scripts and metadata used to construct the panels in Figure 1, please refer to https://github.com/NBChub/meta_data_Figure1.

### Trans-AT PKS BGC mining and analysis

In analysis of trans-AT Polyketide Synthase (PKS) BGCs, we initiated our analysis by clustering all BGC regions identified by antiSMASH using the tool BiG-SCAPE ^36^, with the parameters --cutoff 0.3 --include_singletons --hybrids-off --mibig. This initial clustering utilizes a strict cutoff and does not use the --mix parameters for quick and conservative identification of gene cluster families (GCFs). The nodes in the resulting network were then colored using the predicted compound with highest ranked similarity score against known reference BGCs in the MIBiG database using the ClusterBlast function in antiSMASH. We examined each connected component within the graph, selecting only those containing regions or MIBiG BGCs identified as trans-AT PKSs for further analysis. The BGCs in the filtered subgraph were then re-analyzed using BiG-SCAPE with the parameters set to --mode glocal --include_singletons --mix --clans-off --cutoffs 0.4. The network was visualized using Cytoscape (v3.10.1)^37^ and the results were then manually examined in detail to confirm the presence of a trans-AT PKS producing BGC, based on the domain organization. Subsequently, Clinker (version 0.028)^38^ was used to produce comparison plots to manually validate transAT-PKS classification. This step identified similarities between the trans-AT PKS BGCs confirmed experimentally in the MIBiG secondary metabolite database and the BGC regions in the G1034 dataset. In cases where multiple product classes were present in one region,the known-cluster-blast feature in antiSMASH was employed for manual validation to accurately identify the trans-AT PKS BGC. Moreover, we evaluated the BGCs by comparing their domain organizations to that of the reference BGCs, which were similar and supported by experimental evidence from the literature.

Finally, based on these comprehensive efforts, we categorized the BGCs into three groups: trans-AT PKS BGCs producing a putative known molecule, unknown trans-AT PKS producing BGCs or BGCs discarded as unverified trans-AT PKSs. Analysis of this section can be found in: https://github.com/NBChub/transAT_G1034.

## Results and discussion

The total number of bacterial genome sequences is steadily increasing over the last 15 years. However, the number of complete genomes is still relatively scarce. For the Family *Streptomycetaceae,* as of June 30th 2023, only 501 complete genomes were deposited in the NCBI database (Figure 1A). Here, we present and analyze a dataset which more than doubles this number of publicly available, high-quality, and complete genomes from important genera of filamentous Actinomycetia (Figure 1A and B). We then use the dataset to show how high quality assemblies can be used for analysis which were not previously possible with fewer or more fragmented genomes.

### Sampling and basic statistics on raw data and assemblies

We isolated almost 2000 strains of filamentous Actinomycetia from soil, and sequenced more than 1,000 of them, with ∼90% of them assembling into complete chromosomes. The 1,034 Actinomycetia strains presented here were primarily sampled in Denmark, with 575 strains from Denmark having GPS coordinates, and 155 strains not having recorded GPS coordinates (Figure 2A, map and inner green dots, respectively). The soil samples were taken from all parts of the country of Denmark including islands such as Anholt, Bornholm and Samsø. In addition, there are 304 strains in the dataset which stem from soil samples collected outside of Denmark (Figure 2A, outer green dots).

**Figure 2.**
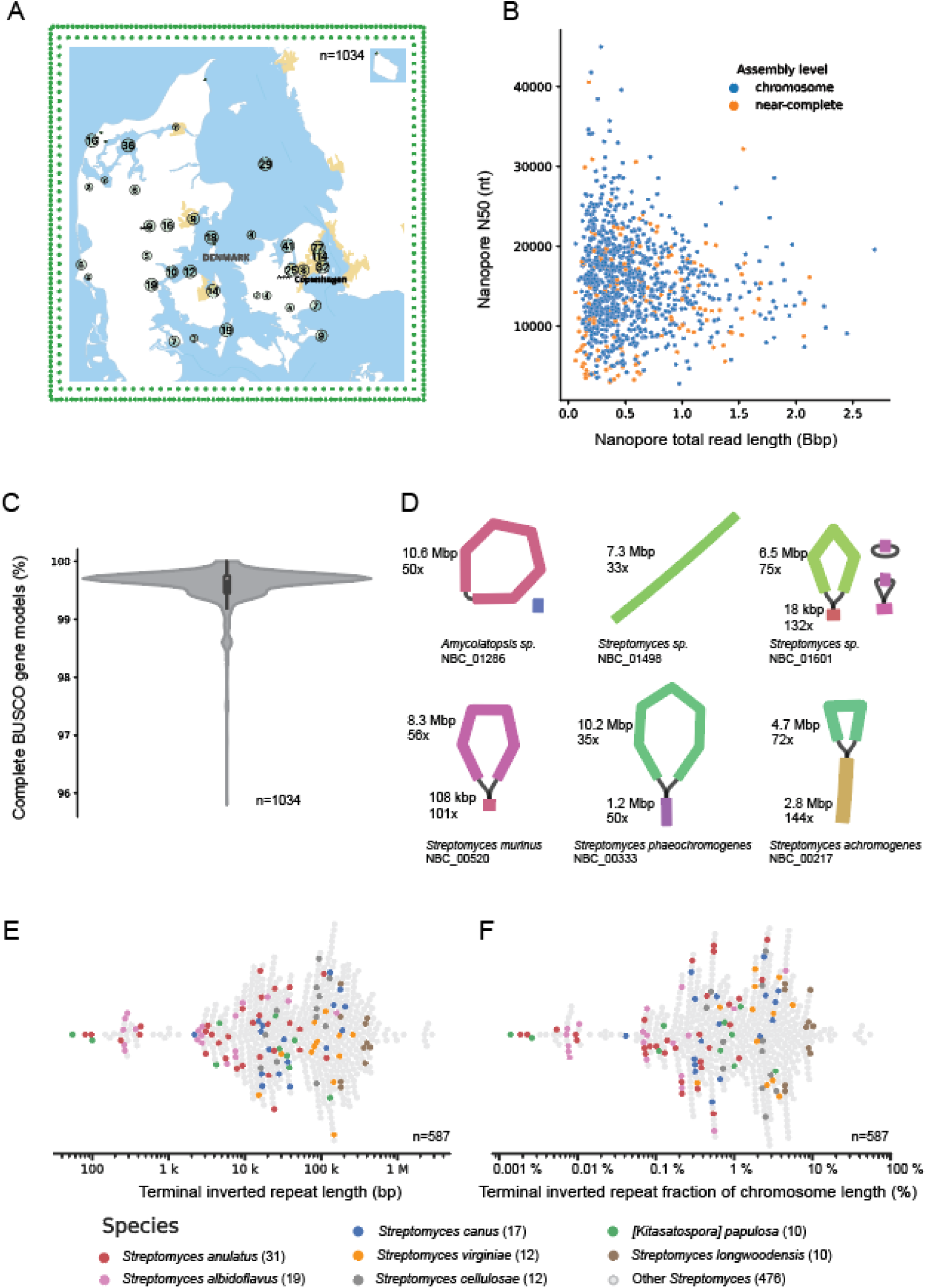
Sampling, genome quality statistics, and topology of genomes. A, map of Denmark showing the sampling locations of the strains presented in this study. The green dots on the inner border show the 155 Danish strains without GPS coordinates, and the 304 dots in the outer border show the strains collected outside of Denmark. B, nanopore data used for assemblies, with N50 and total length of raw nanopore reads shown for chromosome level assemblies (blue) as well as near-complete assemblies (orange). C, Core gene presence measured using the BUSCO actinobacteria_class set of gene models. D, six examples of topologies from circular to linear with increasingly large Terminal inverted repeat. E and F, overview of the size and chromosome fraction taken up by the Terminal Inverted Repeat (TIR). The colored dots correspond to the named species with more than 10 members and show that within species, the TIR size and fraction varies 100 fold.

Two technical factors are crucial for the completeness of a long read genome assembly: the length of reads and the total amount of data. Using an optimized, transposase based nanopore library kit protocol ^18^, we achieved a median nanopore read N50 of 15,376 bp (s.d.= 5,720, n=881) for strains which assembled completely to chromosome level, and a median read N50 of 13,055 bp (s.d. = 6,721 bp, n=153) for strains which did not fully assemble (“WGS” level in NCBI terminology). The median total amount of nanopore data per strain was 427,595,032 bp for chromosome level assemblies (s.d.= 393,977,543 bp, n=881), and 549,676,990 bp (s.d.=469,155,351 bp, n=153) for near-complete assemblies (Figure 2B, each dot represents one strain). The N50 of chromosome level assemblies is longer than that of near complete assemblies (median of 15 kbp vs 12 kbp) which likely reflect that the bottleneck to completely assemble Actinomycetia genomes is the N50, where reads longer than the longest non-terminal repeat is necessary to fully resolve the chromosome organization. The six genomes which did not yield complete assemblies when the N50 was longer than 25 kbp were due to manual filtration as the coverage plots for those strains could not be explained or the issues resolved. The median total nanopore data amount per sample is actually higher in near-complete than in chromosome level assemblies, which reflects extra sequencing effort for strains which did not fully assemble after the initial data generation. However, for some strain and N50 combinations, adding data does not help to resolve a chromosome if the N50 is not sufficiently large. For a comprehensive examination of genome assembly pitfalls, see ^18, 39^.

Each genome was polished with illumina data. The median number of read pairs for the 1,034 illumina datasets are 5,802,158 read pairs (SD 2,191,734, q1 4,580,544, q3 6,971,980 read pairs), which is the rough equivalent of coverage 175 for a 10 Mbp genome. However, for a single genome, the illumina coverage was below 30 x (coverage 8, NBC_00443, illumina read pairs: 246,947). From the Polypolish report we calculated the median number of bases not covered by the illumina reads to be 167 bp or 0.0019 % of the chromosome length (SD 9,709 bp, q1 59 bp, q3 462 bp, uncovered bases on the longest contig in each assembly). However, in a few cases, the number of uncovered bases was larger, up to 208,161 bp(2.2918 %) for the chromosome of strain NBC_01738, which could affect the polishing quality (Supplementary Material S1). A large number of uncovered bases can be caused by poor library quality, which again can be caused by fragmentation, insufficient amount of input DNA, or excessive PCR amplification. Since a median of only a few 100 bp are not covered by illumina data, we consider the quality of the generated illumina data optimal for polishing, however certain types of errors will remain when using short reads for polishing^25^. In line with previous findings on GC bias in illumina data ^17^, more uncovered bases are seen for genomes with higher GC % (Supplementary Material S1).

The genome size for the 1,034 strains range between 2,470,643 bp (*Micrococcus luteus* NBC_00112) and 14,072,637 bp (*Embleya sp.* NBC_0888) with a median genome length of 9,088,824 bp (quantile 1 8,295,182 bp, quantile 3 10,182,039 bp, standard deviation 1,479,678 bp). The GC content in the genomes ranges between 65 % (*Nocardia vinacea* NBC_01743) and 74 % (*Actinomycetospora sp.* NBC_00405), with a median GC content of 71 %. The number of protein coding genes vary from 2,194 genes to 12,441 genes with a median 7,816 genes (n=1034, standard deviation 1263 genes). The number of rRNA operons vary from 2 (e.g. *Kribbella soli* NBC_00256) to 12 (e.g. *Kitasatospora purpeofusca* NBC_01629), though most genomes have around 6 rRNA operons. See Supplementary Material S1 for a comprehensive overview of the basic statistics of the 1,034 genomes including number of tRNAs, rRNAs, the GC-content, genome lengths, BUSCO scores, etc.

A key metric for genome quality is identification of essential, conserved genes, often referred to as core genes. We explored this using the Benchmarking Universal Single-Copy Orthologs (BUSCO) system with the most specific dataset containing all 1,034 genomes: the actinobacteria_class_odb10 dataset of 356 gene models. As seen in Figure 2C, almost all genomes have more than 99 % complete BUSCO genes from the Actinobacteria_class dataset, with a minimum of 95.8 % complete BUSCO gene models (NBC_01737). Though a median of 99.7 % complete BUSCO gene models (SD 0.33 %, q1 99.5 %, q3 99.7 %) can be considered close to perfect, closer inspection of the “fragmented” and “missing” BUSCO genes reveal that out of the 1,466 total missing and fragmented BUSCOs in 1,034 genomes, 886 of them (60 %) have the same BUSCO gene marked as “fragmented” (334658at1760, “cold-shock protein”). This potential artifact particularly affected the genus *Streptomyces*, where 858 out of 886 genomes (97 %) were found to have the gene ‘fragmented’. This could be due to an adaptation in the *Streptomyces* genus, or a technicality in the actinobacteria_odb10 BUSCO dataset. We do not use the BUSCO category “duplicated” as a quality metric, as the validity could be obscured by BGCs carrying resistant versions of the core gene product they are targeting^40^, or simply by non-lethal gene duplications. We did, however, use the duplication score to exclude genomes from the dataset if more than 100 duplicated genes were identified. For three genomes, (NBC_00217, NBC_00887 and NBC_00899), the duplication score was high (16.9 %, 26.7 % and 27.2 %, respectively), and is in all three cases caused by unusually long TIRs duplicating a part of the core genome, possibly as a result of recent and potentially unstable genomic rearrangements.

### Comparative genomics reveals a high taxonomic diversity of the G1034 dataset

In order to estimate the diversity of the G1034 dataset, we performed taxonomic analysis at the whole genome level. GTDB-based taxonomic assignment showed that 1,034 genomes were distributed across 27 genera including *Streptomyces* (n=885), *Micromonospora* (n=36), *Kitasatospora* (n=27), and *Nocardia* (n=21). This study significantly increased the number of complete genomes for several of these genera. One of the strains failed GTDB-Tk run (NBC_0135) and another (NBC_01309) was not assigned to any of the GTDB genera. This strain NBC_01309 with the assigned family Streptomycetaceae represents a potentially novel and unique genus based on genome sequence data. These two strains are not included in the analysis in this section.

We further note that 570 of the 1,032 genomes could be assigned to one of the 145 GTDB-defined species. The remaining 462 genomes were not assigned to any of the taxonomically or computationally defined species. Thus, they likely represent new species that haven’t been described before and thus expands the existing phylogenetic diversity of the *Streptomyces* and other genera. We carried out MASH-based analysis^35^ to detect diversity among these 462 genomes. A whole genome similarity network was reconstructed with 95% similarity as a cutoff for edges. Using a community detection algorithm, we identified 244 communities in the network that can be computationally assigned as different species with 95% similarity cutoff. Some of these MASH-based species had multiple genomes (e.g. 28, 16, 13, 10, etc). As many as 171 of the species were represented by only a single genome, where 58 of the GTDB-assignend species were represented by a single genome. Overall these statistics highlight the diversity of the sequenced collection. See Supplementary material S1 for NCBI, GTDBtk and MASH taxonomic assignment for each strain.

Next, we inferred phylogenetic trees of the G1034 dataset and the dataset of the genus *Streptomyces* with publicly available genome (Figure 1C & D) using GTDB-Tk ^41^. The different species in the G1034 dataset are widely distributed and include representatives of most sub-phyla. The phylogenetic tree of all complete *Streptomyces* genomes further illustrated the diversity of the dataset in the context of existing public data. We observed that there are certain clades of the tree that are overrepresented and very few that do not contain strains from our collection. It is likely that there is still high potential to isolate novel species from this genus and thereby expand the phylogenetic diversity of public databases.

As expected, the G1034 dataset is a rich source of specialized biosynthetic gene clusters. In an analysis of antiSMASH7 annotations, 29,024 regions encoding BGCs were identified in the genomes. As also observed in other studies ^42^, there are clear correlations between genome size and the number of BGCs in the various genera in the dataset (Figure 3A). Many *Actinomycetia* BGCs are hybrids between several biosynthetic types (e.g, type I PKS and NRPS). To generate BGC abundance statistics, we used the antiSMASH proto-clusters^43^, as the protocluster annotation level takes into account the composition of the hybrid BGCs. BGCs coding for the biosynthesis of terpenoids (5866), NRPS (5291) and type I PKSs (4099) are most abundant in the strains, but also siderophores (2396), RiPPs (1986), type III PKS (1848), butyrolactones (1766) and type II PKS (1545) containing BGCs are widespread.

**Figure 3.**
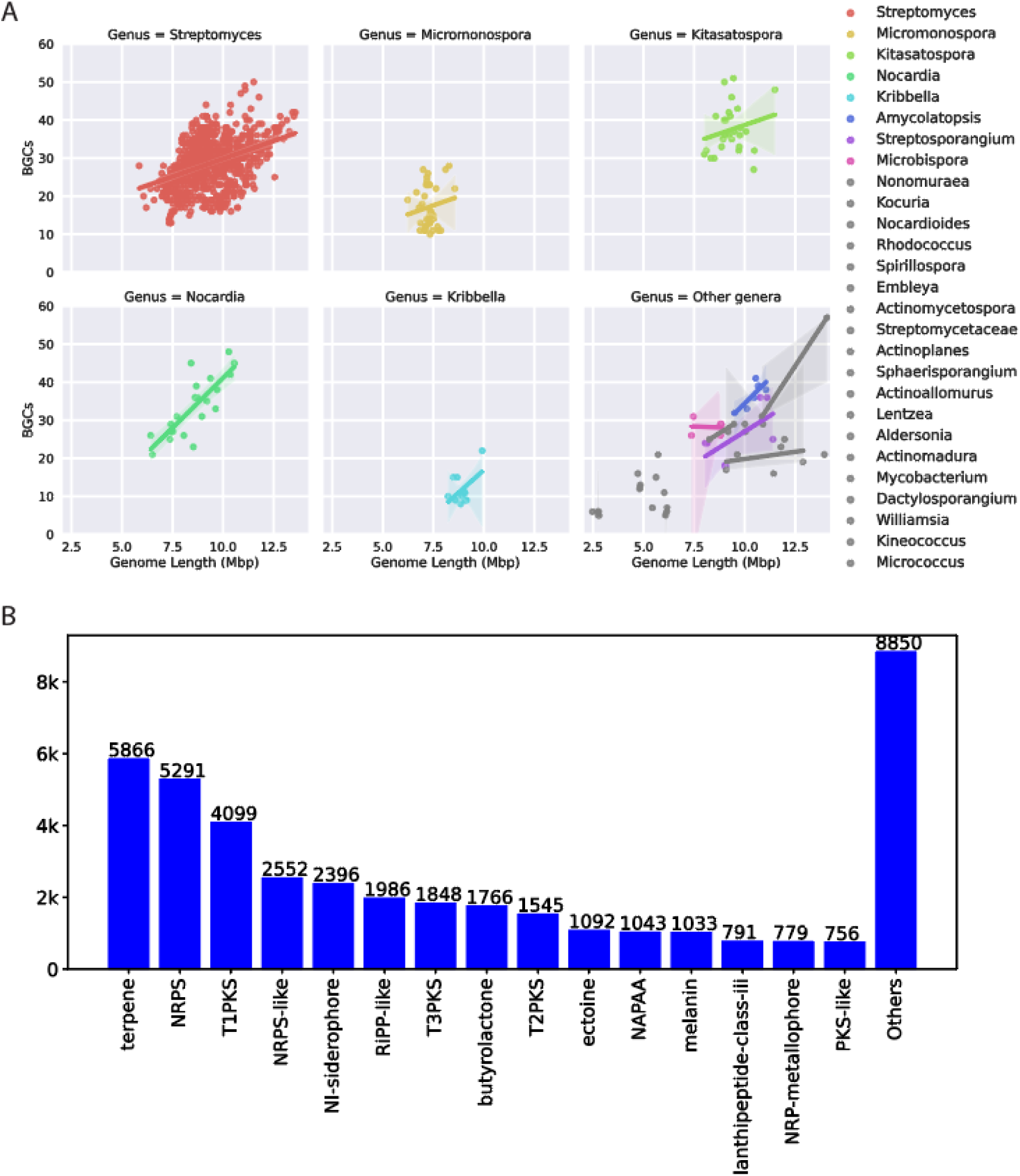
Genome lengths and basic BGC statistics. A) Distribution of number of BGC regions against genome length. Individual scatter plots with regression models represented individually for the top five genera, whereas rest of the genera are combined in the bottom right subpanel. B) Types of BGCs (on antiSMASH protocluster-level) across all 1,034 genomes from NBC dataset distributed across various types. Top 15 types are represented by bars whereas the remaining types are combined in “Others”. Hybrid cluster types are represented in all individual protoclusters in these statistics.

### *Streptomyces* chromosomal organization

While most bacterial genomes are circular, most *Streptomyces* genomes are linear and carry a linear inverted repeat on the ends of the chromosome arms ^44^, overlapping with, but not identical to the chromosome arms. Of the 1,034 genomes in this publication, 886 belong to the genus *Streptomyces*, of which 742 are chromosome level assemblies. We used the assembly repeat graph from Flye ^21^ to manually determine the topology of each individual genome. In Figure 2D, six examples of different chromosome topologies are shown, both circular and linear with increasingly large unresolved Terminal Inverted Repeat from 18kbp to the longest terminal inverted repeat (TIR) in the dataset of 2.8 Mbp. The assembly repeat graphs has been suggested to be a useful guide in correctly assembling a chromosome where the repeats vastly outsize the read length, as is the case for most of the linear inverted repeats of *Streptomyces* (https://github.com/fenderglass/Flye/issues/610#issuecomment-1629027346).

#### *Streptomyces* show strong variability in the length of the terminal inverted repeats

Out of a total of 742 chromosome level *Streptomyces* genomes, TIRs could be identified in 79 % (587/742), while in 21 % (155/742), no TIR was identified using our BLAST based methodology. The strains without identified TIRs could either not contain them, or perhaps the assembly or identification process somehow fell short for these strains, for example the nanopore or illumina polishing could have truncated the extreme chromosome ends. Plotting the TIR lengths both as absolute values and as a fraction of the total chromosome length (including both TIRs in a genome) reveal that the size of the TIRs range from less than 100 bp to almost 3 Mbp, and from less than 0.01 % to more than 50 % of the total chromosome size (Figure 2E and F, see Supplemental material S1 for the data used for the figures). Most of the TIRs are in the 10,000 to 150,000 bp range or 0.3 % to 10 % of the total genome size (median 50,618 bp, q1 14,006 bp, q3 147,244 bp, SD 274,156 bp). The huge size range of more than 1,000-fold showcases the highly dynamic nature of the TIR. We then analyzed the distribution within the named species with more than 10 members (Figure 2E and F, colored dots), and found that even within species, the length of the TIR is highly dynamic ranging for example from few 100 bp to more than 100,000 bp in the most abundant species in the dataset *Streptomyces anulatus* (Figure 2E and F, red dots). This dynamic TIR correlates to observations of *Streptomyces* strains undergoing large changes in the TIR during growth in a laboratory, and to frequent reports of the loss of one or both chromosome arms due to double strand breaks ^11, 44^. Understanding these processes also has very important implications for engineering *Streptomyces*, i.e. in knockout experiments, where a duplicate gene can be harder to modify or in the worst case even can cause megabase-sized deletions. Since BGCs are commonly examined and often found on TIRs, there is a high risk of misinterpreting experimental results if the TIR is not correctly accounted for.

#### *Streptomyces* show conserved core gene placement

In order to explore the genome organization of the 742 linear *Streptomyces* genomes, we first oriented all genomes according to the direction of the *dnaA* gene. For circular genomes, this gene is commonly used to rotate and orient the genome sequence ^45^. For four strains, there were two annotated *dnaA* genes, however in all cases the *dnaA* genes were located on the same strand. Around 50 % of genomes were flipped, as would be expected from a random initial genomic direction. We used the 124 HMM gene models in the bacteria_odb10 (v.2020-03-06) BUSCO dataset to identify single copy essential genes conserved in all Bacteria, as these can be considered the most conserved core genes. The position of each gene was then extracted from the genbank files and used for the probability density estimates in Figure 4A and B, with the middle of gene coordinates being used for calculations. This revealed that the placement of the core genes in linear *Streptomyces* genomes is extremely structured: the *dnaA* gene is the most centrally placed conserved gene in the complete *Streptomyces* genomes, encoding the DNA replication initiation protein. It is located exclusively near the mathematical center of the linear chromosome with a median distance of −21,173 bp, just upstream of the chromosome center (q1: −130,158 bp, q3: 77,679 bp, SD: 214,661 bp). An additional 11 genes are consistently located around the chromosome center, among them genes encoding ribosomal proteins and tRNA ligases (Figure 4A, red and orange hues).

**Figure 4.**
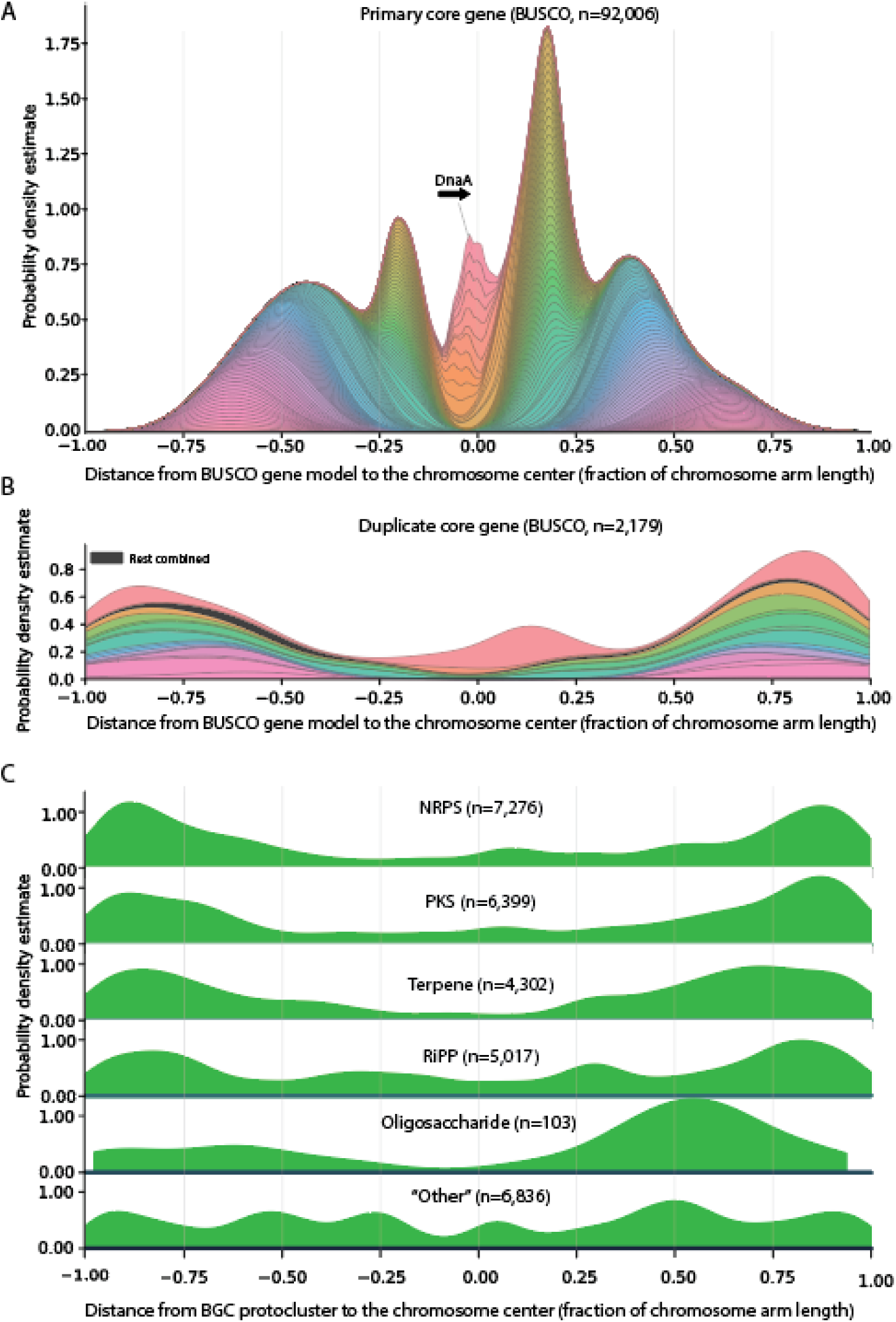
Distribution of core genes and BGC categories along the linear *Streptomyces* chromosome. Kernel density plots of core genes and BGC protoclusters along the 742 complete linear *Streptomyces* genomes in the present study. A, placement of 124 bacterial core genes from the BUSCO Bacteria dataset, showing central and highly organized placement in relation to the chromosome center. B, secondary copy core genes showing a distal placement for all but one core gene. C, the six antiSMASH categories of protoclusters, showing a distal placement for all except for the ‘other’ category.

Generally, all 124 bacterial core gene models seem to have a preferred distance to the chromosome center (see Supplementary Material S4 for complete legend for Figure 4A. See Supplementary Material S5 for the tables used for Figure 4). From the small ribosomal subunit, 10 out of 17 genes are in the chromosome arms 12-25 % of the arm length distance from the chromosome center (Figure 4A, green and yellow hues) in a clear focal point likely reflecting a cluster of ribosomal operons. In the same focal point, 14 out of 25 genes from the large ribosomal subunit are found. Other ribosomal proteins such as L20, L35, and S4 are positioned halfway towards the chromosome ends (Figure 4A, purples). Generally, only a few genes shared by all of Bacteria are placed outside of the inner 75 % of the chromosome, which can be considered the central genomic region. For genes unaffected by the direction of replication, the density on either side of the chromosome center should be of equal size. However, there seem to exist a preferred position of a group of genes relating to the direction of the *dnaA* gene, where several of the ribosomal genes are more likely to be found downstream than upstream of *dnaA*. This effect can be observed for most core genes, but it seems that the effect decreases with the distance to the chromosome center. This observation of core gene placement is in agreement with previous observations on a smaller *Streptomyces* dataset ^15, 46^ and several genome organization descriptions ^47, 48^, where core genes were found close to the central chromosome in a ‘central compartment’.

In most genomes, a few BUSCO genes are found in duplicate. A total of 2,179 genes were found in duplicate from 742 complete *Streptomyces* genomes, giving an average duplicate count from the bacterial BUSCO dataset of approximately 3 genes per genome (Figure 4B). Most of the duplicated genes are positioned away from the chromosome center with a single prominent exception of the ribosomal protein S18 gene (1940575at2), which have densities both in the central chromosome and on the chromosome arms. For the remaining secondary duplicate genes, the density estimates are shifted towards the ends of the chromosome arms, away from the chromosome center. This observation is in line with the observation that BGCs often carry a resistant version of the target of the specialized metabolite^40^. The observations on duplicate core genes are weakened by the methodology: the primary and duplicate gene copies are not distinguished on the base of sequence identity or divergence, but on distance to the chromosome center, forcing the “secondary” copy to be more distal than the “primary” copy.

#### BGC are not placed randomly on linear *Streptomyces* genomes

Having documented the conserved placement of core genes along the linear *Streptomyces* chromosome, we next investigated the placement of BGCs. The observation that biosynthetic gene clusters often are located towards the linear chromosome ends has been suggested since the first complete *Streptomyces* genome sequences became available^14^. antiSMASH (v.7.0.0) operates with 81 rules, each describing the core biosynthetic requirements of a specific type of BGC, e.g. Type 1 Polyketide Synthase (T1PKS) or lassopeptide. Each cluster type has a predefined neighborhood on either side of the core cluster (e.g. 20 kbp on either side of T1PKS core genes), which together constitutes a protocluster. Overlapping protoclusters are annotated as a ‘region’, the primary output from the antiSMASH software. Each protocluster is annotated as one of six categories, responsible for production of the most common groups of compounds: non-ribosomal peptide synthase (NRPS), polyketide synthase (PKS), terpenes, ribosomally synthesized and post-translationally modified peptides (RiPP), Oligosaccharide, and ‘other’. Depending on the purpose of the analysis, we either analyze protocluster or region in this paper. For a comprehensive overview of the annotation layers generated by antiSMASH, please refer to ^6, 43^. For an overview of the protocluster types, categories, and counts, please refer to Supplementary Material S6.

The global distribution of BGCs on linear *Streptomyces* chromosomes was explored using the six antiSMASH categories (Figure 4C). In all cases except for “other”, the BGC category is preferentially located towards the end of the chromosome, confirming previous observations that most specialized metabolite clusters are placed distally on the linear *Streptomyces* chromosome. The “other” category includes a diverse range of BGCs, which could explain the lack of a clear pattern in the distribution. For the category oligosaccharide, the density is skewed to the right arm, however this could be an artifact from a low number of protocluster observations and an uneven taxonomic distribution (n=103). In total, this analysis is based on 29,933 Streptomyces protoclusters. We chose to represent the annotation level “protocluster” to get a comprehensive overview of BGCs not impacted by adjacent BGCs which antiSMASH did not resolve. However, since some clusters are true hybrids consisting of more than one type of protocluster, this approach risks overestimating the total number of clusters in a dataset. Conversely, had we used the antiSMASH annotation level ‘region’, the number of common cluster types would certainly be underestimated, as clusters physically near –but unrelated to– other clusters would be in the same region and thus only be represented once, severely reducing the dataset near the linear chromosome ends.

We then proceeded to analyze the placement of a few common antiSMASH cluster types, (T1PKS, T2PKS, T3PKS, trans-AT PKS), or cluster types which had a clear distribution along the linear *Streptomyces* chromosomes (NI-siderophore, NRP-metallophore, Ectoine, Melanin, Arylpolyene). Plots for the remaining cluster types (with enough observations for a density plot) can be found in Supplementary material S7.

#### Polyketides

For each of the four PKS types in Figure 5, there is a mostly symmetrical distal distribution, with very few protoclusters found in the central region of the genome. Trans-AT PKSs seem to be heavily skewed to the “right” chromosome arm, however the number of observations is low (139) which means that the distribution could be skewed by a few common cluster families.

**Figure 5.**
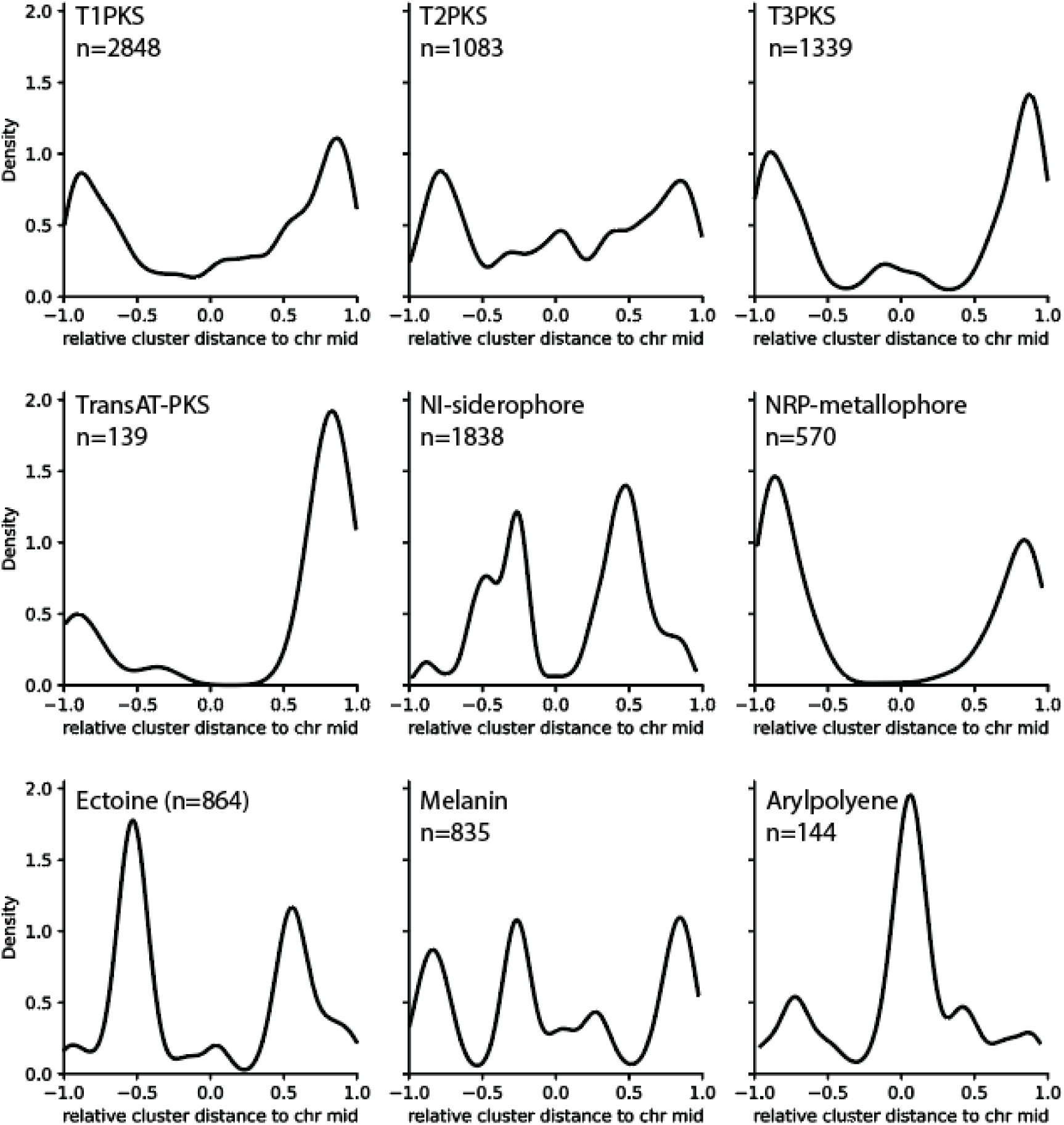
Distribution of select individual BGC types along the linear Streptomyces Chromosome. kernel density plots showing the placement of specific antiSMASH7 protocluster types along the linear *Streptomyces* chromosomes. While the PKS type clusters are distally placed, other types such as NI-siderophore and Arylpolyene are centrally placed.

#### Metallophores

Metal ions, especially Fe(II)/Fe(III), are essential cofactors for many cell functions, and organisms across the whole tree of life possess highly selective uptake systems to acquire these ions from the environment, frequently utilizing chelator-based sequestration. In bacteria, biosynthesis of these metallophores can be distinguished to be either dependent or independent of an NRPS (for an overview, see ^49^). In *Streptomyces* genomes both the NRPS dependent type (called NRP-metallophore, Figure 5, middle right) and the NRPS independent type (called NI-siderophore, Figure 5, center) can be found. Notably, NI-siderophores tend to be placed more centrally on the chromosomes whereas NRP-metallophore clusters occur more distally.

#### Ectoines

Ectoine (Figure 5, lower left) is a compound which plays roles in protecting cells against osmotic stress. The BGCs responsible for production of Ectoine have a mostly symmetric distribution approximately halfway between the linear chromosome center and the chromosome ends, in an area where core genes are often found. This very localized placement suggests a conservation in the distribution along the linear *Streptomyces* chromosome.

#### Melanines

Melanin-type protoclusters (Figure 5, lower middle) are placed all the way along the linear chromosome, with peaks both in the distal parts and the more central parts. Melanin-type clusters are responsible for production of dark or colorful pigment compounds, with several subtypes performing different roles. The best known role for a melanin compound is UV protection in both eu- and prokaryotes, but some melanin-type clusters encode compounds with antibacterial and antiviral activity ^50^, hinting that the distribution of melanins along the linear chromosome could be more located if the BGC type was broken further down into cluster families.

#### Arylpolyenes

Arylpolyenes (Figure 5, lower right), which are pigments with functions similar to carotenoides, is encoded by a special subtype of T2PKS type BGCs ^51^. Arylpolyene BGCs have a prominently central location on the linear chromosome, potentially reflecting a role in primary metabolism.

For the BGC types CDPS, NAPAA, NRPS, NRPS-like, hlgE-KS, terpene, lanthipeptide-class-iv, linaridine, redox-cofactor, RiPP-like, amglyccycl, aminopolycarboxylic acid, betalactone, blactam, indole, nucleoside, and “other”, a distal distribution is also seen, whereas for the remaining types, the distribution is either central or spread out over the whole genome (See Supplemental Information S4 for distribution plots of all cluster types with >50 observations).

### A treasure trove for specialized metabolite biosynthetic gene clusters using trans-AT PKS BGCs as an example

While statistics on the BGC-content give an excellent overview of the biosynthetic potential of the individual strains, they don’t indicate the diversity of BGCs across the dataset. To analyze the biosynthetic potential of a dataset, BGCs can be assigned to gene cluster families (GCFs). GFCs contain similar BGCs that most likely are involved in the biosynthesis of identical or chemically closely related natural products. A common approach for GCF identification includes reconstruction of a similarity metric between BGCs and then grouping BGCs into GCFs based on clustering of the similarity networks. For example, BiG-SCAPE generates BGC-class specific combined distance metric based on similarity metrics^36^. Inclusion of BGCs for known natural products from the MIBiG reference database allows the assessment if the identified GCFs contain BGCs involved in the biosynthesis of known molecules or if they code for potentially novel pathways^52^. A similarity network analysis with BiG-SCAPE indicated that the 29,024 BGC regions of G1034 can be assigned to as many as 7,193 GCFs. 190 of these could be associated with a BGC in the MIBiG reference dataset and thus likely code for known compounds or derivatives of these. We note that 4,400 of the BIG-SCAPE GCFs are unique to individual isolates in the dataset. As the diversity of BGCs and GCFs in the G1034 dataset is enormous, we chose to focus on the BGC type trans-AT PKS as an example of how to query the dataset.

Trans-acyltransferase polyketides constitute a highly diverse group of bacterial multimodular specialized metabolites. These compounds are biosynthesized through diverse biosynthetic pathways, posing significant challenges for their characterization using computational prediction tools. Unlike cis-acyltransferase polyketide synthases (cis-AT PKSs), where acyltransferase domains are integrated within the modules, trans-AT PKSs are characterized by the fact that the acyltransferase is encoded as a separate gene. The polyketide product often diverges from this expected module-based structure, however, common traits of trans-AT PKSs have aided the prediction of these intricate pathways^53^. One example is the catalytic ketosynthase (KS) domains in the trans-AT PKS pathway, which can be grouped into clades correlating with distinct substrate specificity^53, 54^. For a detailed description of the trans-AT PKS biosynthesis see ^55^. These conserved features make detection of trans-AT PKSs possible, however, subsequent manual curation of the BGCs is beneficial when comparing novel BGCs as they may be assigned to a MIBiG reference because they share genes adjacent to the biosynthetic core but not necessarily synthesize similar molecules.

In our analysis of the G1034 dataset, we identified 247 trans-AT hits matching the antiSMASH trans-AT classification rule. Of these, 122 BGCs showed high similarity to 14 BGCs of known compounds. This similarity was confirmed through manual inspection of each BGC by identifying the described core genes in the biosynthesis pathway, as detailed in the methods section. Remarkably, out of the 17 described trans-AT PKS reference BGCs found in the MIBiG database from the phylum Actinomycetia, 14 are found in the G1034 dataset. The 247 transAT-PKS BGCs stem from 87 different species which are represented with fewer than ten trans-AT PKS BGCs, and two species, *Streptomyces* sp003846175 with 26 trans-AT PKS BGCs and *Streptomyces anulatus* with 56 transAT-PKS BGCs. These two species are also among the most well represented in the G1034 dataset with 26 and 25 strains, respectively.

Furthermore, our analysis revealed 41 trans-AT PKS BGCs that did not resemble a MIBiG reference with similar biosynthetic genes. These BGCs were categorized into 7 gene cluster families and 14 singletons with unique domain organization. Given their distinct profiles, these GCFs are presumed to be putative novel BGCs, potentially representing new natural products (see Figure 6B). The remaining 84 BGCs were not included in this analysis as they were found to be part of larger hybrid cluster regions, which if included would interfere with GCF assignment.

**Figure 6:**
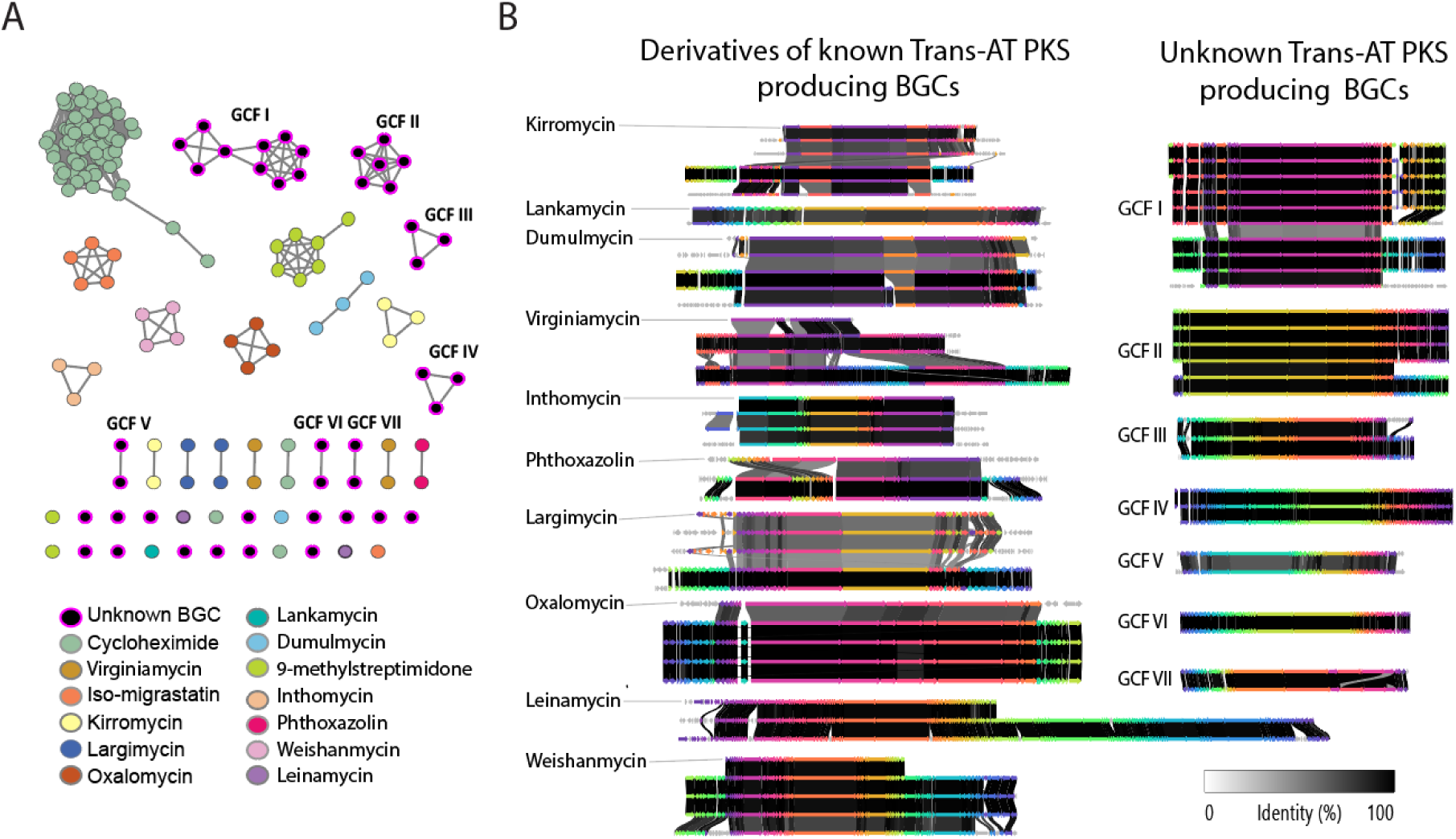
GCF network and BGC region comparison of selected identified GCFs. A, network representation of 163 trans-AT PKS BGCs generated with BIG-SCAPE and positioned by hierarchical clustering. Nodes are colored by putative production of a known trans-AT PKS or described as an unknown trans-AT PKS. B, Clinker gene comparison plots displaying trans-AT PKS BGC regions with similar domain structure as experimentally validated BGCs from the MIBiG secondary metabolite database. The MIBiG entry is on top in each plot. Furthermore, gene comparison plots displaying GCF of trans-AT PKS BGCs with no similar reference are shown on the right. Genome accession numbers and the full network are available in the Supplementary material S8.

In the network representation of likely trans-AT PKS producing BGCs (Figure 6A) the similarity cutoff 0.40 was chosen to best describe the relationship between BGCs. This cutoff highlighted substantial similarities among the BGCs, despite divergent features primarily due to neighboring genes to the biosynthetic core causing the BGCs to deviate from the main network cluster with BGCs that share biosynthetic core properties. A notable example of this scenario are the 5 kirromycin-like BGCs, which, despite a gene rearrangement and distinct flanking regions between its BGCs, share similar biosynthetic core regions, but are split into two network clusters (Figure 6B, yellow). Accession numbers can be found in Supplementary Material S8 and the manually curated overview of trans-AT PKS BGCs can be found in Supplementary Material S9.

### Conclusions

We have collected, sequenced and analyzed 1,034 genomes from filamentous actinomycetes, in a dataset which more than doubles the available HQ complete genomes from the important genus *Streptomyces*. In this dataset, we identified more than 400 genomes from potential new species, as well as genomes from 145 existing species. We here provide the first large scale analysis of the terminal inverted repeats of *Streptomyces* genomes. We analyzed the placement of core genes and BGCs along the linear *Streptomyces* genome and found a conserved pattern, where features are not only located within the genomics compartments, but each feature is also organized with a certain distance to the chromosome center. Finally, we investigated trans-AT PKSs as an example of the BGC-focus analysis possible, and found that 14 out of 17 known actinomyces derived trans-AT PKS’s could be identified in the dataset, along with several new trans-AT PKS cluster families. With this study, we show the possible analysis which can only be performed on chromosome resolved complete genomes, and provide a valuable source of genomic information to be analyzed in future studies.

## Supporting information

Supplementary_material_S1_Statistics_table_per_strain

Supplementary_material_S2_coverage_plots

Supplementary_material_S3_list_of_countries

Supplementary_material_S4_core_gene_placement_legend

Supplementary_material_S5_Tables_for_fig_4_placement

Supplementary_material_S6_Table_of_antiSMASH_cluster_types_to_categories_and_occurences

Supplementary_material_S7_Antismash_cluster_type_plots_all_protocluster_types

Supplementary_material_S8_figure_clinkers_w_names

Supplementary_material_S9_tables_for_transAT-PKS_analyses

## Data availability

All sequence data was deposited at NCBI under BioProject (PRJNA747871). NCBI-Accession numbers for the 1034 individual strains are listed in the BioProject entry and in Table S1. See Materials and Methods for links to the software used to analyze the data.

## Acknowledgements

We thank Oliwia Vuksanovic and Alexandra Hoffmeyer for excellent technical assistance and Jens Christian von der Maase for permission to collect soil samples in Ørkenen, Anholt, Denmark. We thank the NCBI GenBank Direct Submission team and particularly Leigh A. Riley and Linda Frisse for guidance in the process of submitting Biosamples, reads, and assemblies.

This work was funded by grants from the Novo Nordisk Foundation [NNF20CC0035580, NNF16OC0021746].

## Supplemental information

| New numbering | Line of first mention |
| --- | --- |
| Supplementary_material_S1_Statistics_table_per_strain | l109 |
| Supplementary_material_S2_coverage_plots | l172 |
| Supplementary_material_S3_list_of_countries | l277 |
| Supplementary_material_S4_core_gene_placement_legend | 539 |
| Supplementary_material_S5_Tables_for_figure_4_placement | l540 |
| Supplementary_material_S6_Table_of_antiSMASH_cluster_types_to_categories_and_occurrences | l596 |
| Supplementary_material_S7_Antismash_cluster_type_plots_all_protocluster_types | l618 |
| Supplementary_material_S8_figure_clinkers_w_names | l737 |
| Supplementary_material_S9_tables_for_transAT-PKS_analyses | l738 |

Potential fig for graphical abstract: all core genes and all BGCs from complete *streptomyces* genomes.

## Notes

### Competing Interest Statement

The authors have declared no competing interest.

### Summary of Updates

Abstract had a confusing/not correct wording. The sentence now reads "Filamentous Actinobacteria, recently renamed Actinomycetia, are the most prolific source of microbial bioactive natural products."

## References

1. Barka Essaid Ait et al. Taxonomy, Physiology, and Natural Products of Actinobacteria. Microbiol. Mol. Biol. Rev. 80, 1–43 (2015).

2. Perry, R. P. & Kelley, D. E. Inhibition of RNA synthesis by actinomycin D: characteristic dose-response of different RNA species. J. Cell. Physiol. 76, 127–139 (1970).

3. Beck, C. et al. Activation and Identification of a Griseusin Cluster in Streptomyces sp. CA-256286 by Employing Transcriptional Regulators and Multi-Omics Methods. Molecules 26, (2021).

4. Crits-Christoph, A., Bhattacharya, N., Olm, M. R., Song, Y. S. & Banfield, J. F. Transporter genes in biosynthetic gene clusters predict metabolite characteristics and siderophore activity. Genome Res. 31, 239–250 (2020).

5. Alanjary, M. et al. The Antibiotic Resistant Target Seeker (ARTS), an exploration engine for antibiotic cluster prioritization and novel drug target discovery. Nucleic Acids Res. 45, W42–W48 (2017).

6. Blin, K. et al. antiSMASH 7.0: new and improved predictions for detection, regulation, chemical structures and visualisation. Nucleic Acids Res. (2023) doi:10.1093/nar/gkad344.

7. Blin, K., Kim, H. U., Medema, M. H. & Weber, T. Recent development of antiSMASH and other computational approaches to mine secondary metabolite biosynthetic gene clusters. Brief. Bioinform. 20, 1103–1113 (2019).

8. Baltz, R. H. Genome mining for drug discovery: progress at the front end. J. Ind. Microbiol. Biotechnol. 48, (2021).

9. Carretero-Molina, D. et al. Discovery of gargantulides B and C, new 52-membered macrolactones from Amycolatopsis sp. Complete absolute stereochemistry of the gargantulide family. Organic Chemistry Frontiers 9, 462–470 (2022).

10. Sánchez-Navarro, R., et al. Long-Read Metagenome-Assembled Genomes Improve Identification of Novel Complete Biosynthetic Gene Clusters in a Complex Microbial Activated Sludge Ecosystem. mSystems 7, e0063222 (2022).

11. Tidjani, A.-R., Bontemps, C. & Leblond, P. Telomeric and sub-telomeric regions undergo rapid turnover within a Streptomyces population. Sci. Rep. 10, 7720 (2020).

12. Aigle, B. et al. Genome mining of Streptomyces ambofaciens. J. Ind. Microbiol. Biotechnol. 41, 251–263 (2014).

13. Yang, C.-C., Tseng, S.-M. & Chen, C. W. Telomere-associated proteins add deoxynucleotides to terminal proteins during replication of the telomeres of linear chromosomes and plasmids in Streptomyces. Nucleic Acids Res. 43, 6373–6383 (2015).

14. Bentley, S. D. et al. Complete genome sequence of the model actinomycete Streptomyces coelicolor A3(2). Nature 417, 141–147 (2002).

15. Kim, J.-N. et al. Comparative Genomics Reveals the Core and Accessory Genomes of Streptomyces Species. J. Microbiol. Biotechnol. 25, 1599–1605 (2015).

16. Lorenzi, J.-N. et al. Ribosomal RNA operons define a central functional compartment in the Streptomyces chromosome. Nucleic Acids Res. 50, 11654–11669 (2022).

17. Browne, P. D. et al. GC bias affects genomic and metagenomic reconstructions, underrepresenting GC-poor organisms. Gigascience 9, (2020).

18. Alvarez-Arevalo, M. et al. Extraction and Oxford Nanopore sequencing of genomic DNA from filamentous Actinobacteria. STAR Protoc 4, 101955 (2022).

19. Seshadri, R. et al. Expanding the genomic encyclopedia of with 824 isolate reference genomes. Cell Genom 2, 100213 (2022).

20. Krueger, F. Trim Galore!: A wrapper around Cutadapt and FastQC to consistently apply adapter and quality trimming to FastQ files, with extra functionality for RRBS data. (2015).

21. Kolmogorov, M., Yuan, J., Lin, Y. & Pevzner, P. A. Assembly of long, error-prone reads using repeat graphs. Nat. Biotechnol. 37, 540–546 (2019).

22. Wick, R. R., Schultz, M. B., Zobel, J. & Holt, K. E. Bandage: interactive visualization of de novo genome assemblies. Bioinformatics 31, 3350–3352 (2015).

23. Langmead, B. & Salzberg, S. L. Fast gapped-read alignment with Bowtie 2. Nat. Methods 9, 357–359 (2012).

24. Li, H. Minimap2: pairwise alignment for nucleotide sequences. Bioinformatics 34, 3094–3100 (2018).

25. Wick, R. R. & Holt, K. E. Polypolish: Short-read polishing of long-read bacterial genome assemblies. PLoS Comput. Biol. 18, e1009802 (2022).

26. Zimin, A. V. et al. The MaSuRCA genome assembler. Bioinformatics 29, 2669–2677 (2013).

27. Chaumeil, P.-A., Mussig, A. J., Hugenholtz, P. & Parks, D. H. GTDB-Tk v2: memory friendly classification with the genome taxonomy database. Bioinformatics 38, 5315–5316 (2022).

28. Simão, F. A., Waterhouse, R. M., Ioannidis, P., Kriventseva, E. V. & Zdobnov, E. M. BUSCO: assessing genome assembly and annotation completeness with single-copy orthologs. Bioinformatics 31, 3210–3212 (2015).

29. Jørgensen, T. S., Hansen, M. A., Xu, Z. & Tabak, M. A. Plasmids, viruses, and other circular elements in rat gut. bioRxiv (2017).

30. Caro, H., et al. BioConvert: a comprehensive format converter for life sciences. NAR Genom Bioinform 5, lqad074 (2023).

31. Altschul, S. F., Gish, W., Miller, W., Myers, E. W. & Lipman, D. J. Basic local alignment search tool. J. Mol. Biol. 215, 403–410 (1990).

32. datamash - GNU Project - Free Software Foundation. https://www.gnu.org/software/datamash/.

33. Gurevich, A., Saveliev, V., Vyahhi, N. & Tesler, G. QUAST: quality assessment tool for genome assemblies. Bioinformatics 29, 1072–1075 (2013).

34. Parks, D. H. et al. GTDB: an ongoing census of bacterial and archaeal diversity through a phylogenetically consistent, rank normalized and complete genome-based taxonomy. Nucleic Acids Res. 50, D785–D794 (2022).

35. Ondov, B. D. et al. Mash: fast genome and metagenome distance estimation using MinHash. Genome Biol. 17, 132 (2016).

36. Navarro-Muñoz, J. C. et al. A computational framework to explore large-scale biosynthetic diversity. Nat. Chem. Biol. 16, 60–68 (2020).

37. Shannon, P. et al. Cytoscape: a software environment for integrated models of biomolecular interaction networks. Genome Res. 13, 2498–2504 (2003).

38. Gilchrist, C. L. M. & Chooi, Y.-H. clinker & clustermap.js: automatic generation of gene cluster comparison figures. Bioinformatics 37, 2473–2475 (2021).

39. Wick, R. R., Judd, L. M. & Holt, K. E. Assembling the perfect bacterial genome using Oxford Nanopore and Illumina sequencing. PLoS Comput. Biol. 19, e1010905 (2023).

40. Mungan, M. D. et al. ARTS 2.0: feature updates and expansion of the Antibiotic Resistant Target Seeker for comparative genome mining. Nucleic Acids Res. 48, W546–W552 (2020).

41. Alanjary, M., Steinke, K. & Ziemert, N. AutoMLST: an automated web server for generating multi-locus species trees highlighting natural product potential. Nucleic Acids Res. 47, W276–W282 (2019).

42. Baltz, R. H. Gifted microbes for genome mining and natural product discovery. J. Ind. Microbiol. Biotechnol. 44, 573–588 (2017).

43. Blin, K. et al. antiSMASH 5.0: updates to the secondary metabolite genome mining pipeline. Nucleic Acids Res. 47, W81–W87 (2019).

44. Hoff, G., Bertrand, C., Piotrowski, E., Thibessard, A. & Leblond, P. Genome plasticity is governed by double strand break DNA repair in Streptomyces. Sci. Rep. 8, 5272 (2018).

45. Hunt, M. et al. Circlator: automated circularization of genome assemblies using long sequencing reads. Genome Biol. 16, 294 (2015).

46. Chung, Y.-H. et al. Comparative Genomics Reveals a Remarkable Biosynthetic Potential of the Streptomyces Phylogenetic Lineage Associated with Rugose-Ornamented Spores. mSystems 6, e0048921 (2021).

47. Ikeda, H. et al. Complete genome sequence and comparative analysis of the industrial microorganism Streptomyces avermitilis. Nat. Biotechnol. 21, 526–531 (2003).

48. Weaver, D. et al. Genome plasticity in Streptomyces: identification of 1 Mb TIRs in the S. coelicolor A3(2) chromosome. Mol. Microbiol. 51, 1535–1550 (2004).

49. Miethke, M. & Marahiel, M. A. Siderophore-based iron acquisition and pathogen control. Microbiol. Mol. Biol. Rev. 71, 413–451 (2007).

50. El-Naggar, N. E.-A. & El-Ewasy, S. M. Bioproduction, characterization, anticancer and antioxidant activities of extracellular melanin pigment produced by newly isolated microbial cell factories Streptomyces glaucescens NEAE-H. Sci. Rep. 7, 42129 (2017).

51. Schöner, T. A. et al. Aryl Polyenes, a Highly Abundant Class of Bacterial Natural Products, Are Functionally Related to Antioxidative Carotenoids. Chembiochem 17, 247–253 (2016).

52. Terlouw, B. R. et al. MIBiG 3.0: a community-driven effort to annotate experimentally validated biosynthetic gene clusters. Nucleic Acids Res. 51, D603–D610 (2023).

53. Helfrich, E. J. N. et al. Automated structure prediction of trans-acyltransferase polyketide synthase products. Nat. Chem. Biol. 15, 813–821 (2019).

54. Nguyen, T. et al. Exploiting the mosaic structure of trans-acyltransferase polyketide synthases for natural product discovery and pathway dissection. Nat. Biotechnol. 26, 225–233 (2008).

55. Helfrich, E. J. N. & Piel, J. Biosynthesis of polyketides by trans-AT polyketide synthases. Nat. Prod. Rep. 33, 231–316 (2016).

