## Supplementary figures and images for "A treasure trove of 1,034 actinomycete genomes"

### Supplementary_material_S2_coverage_plots

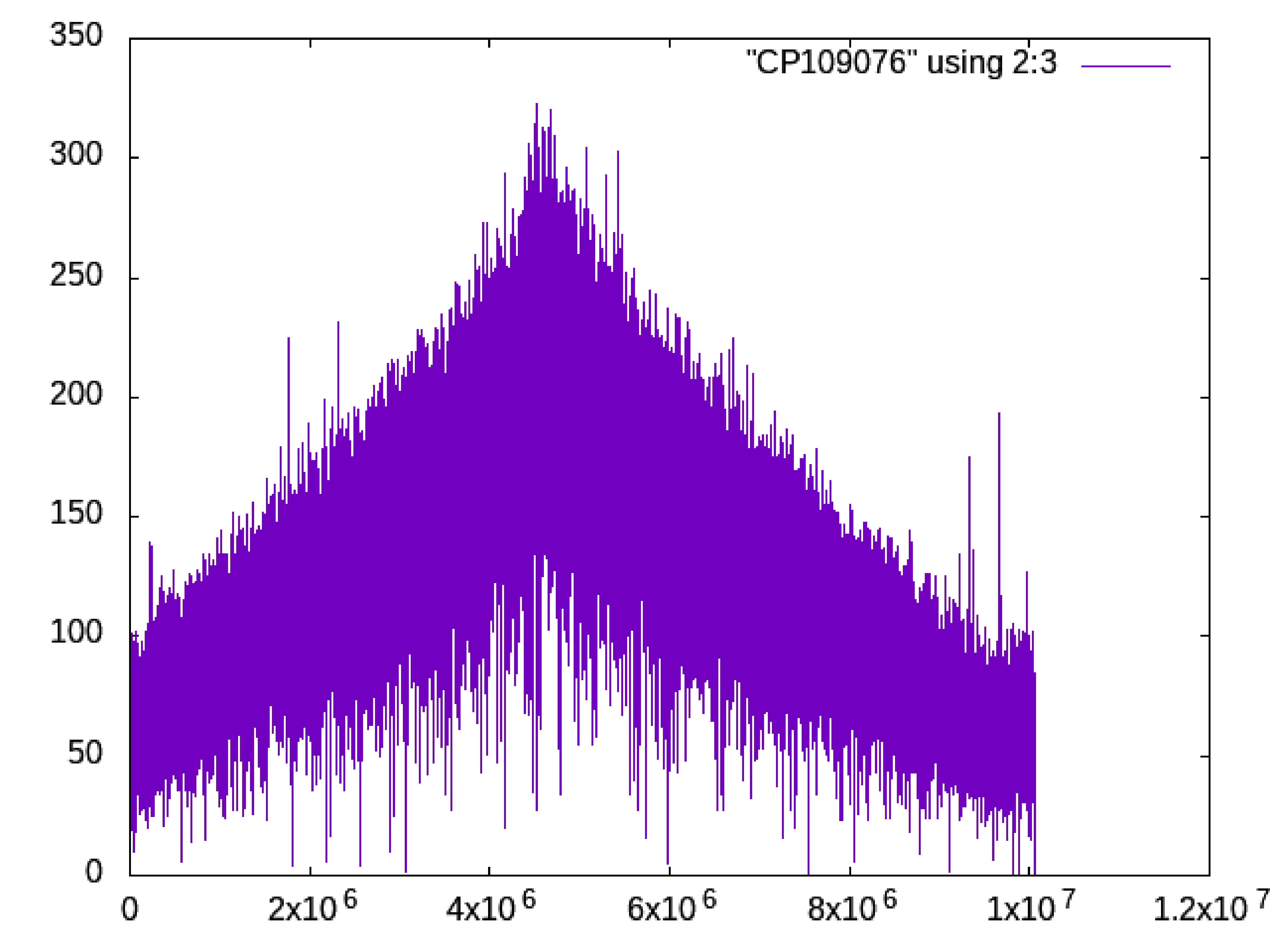

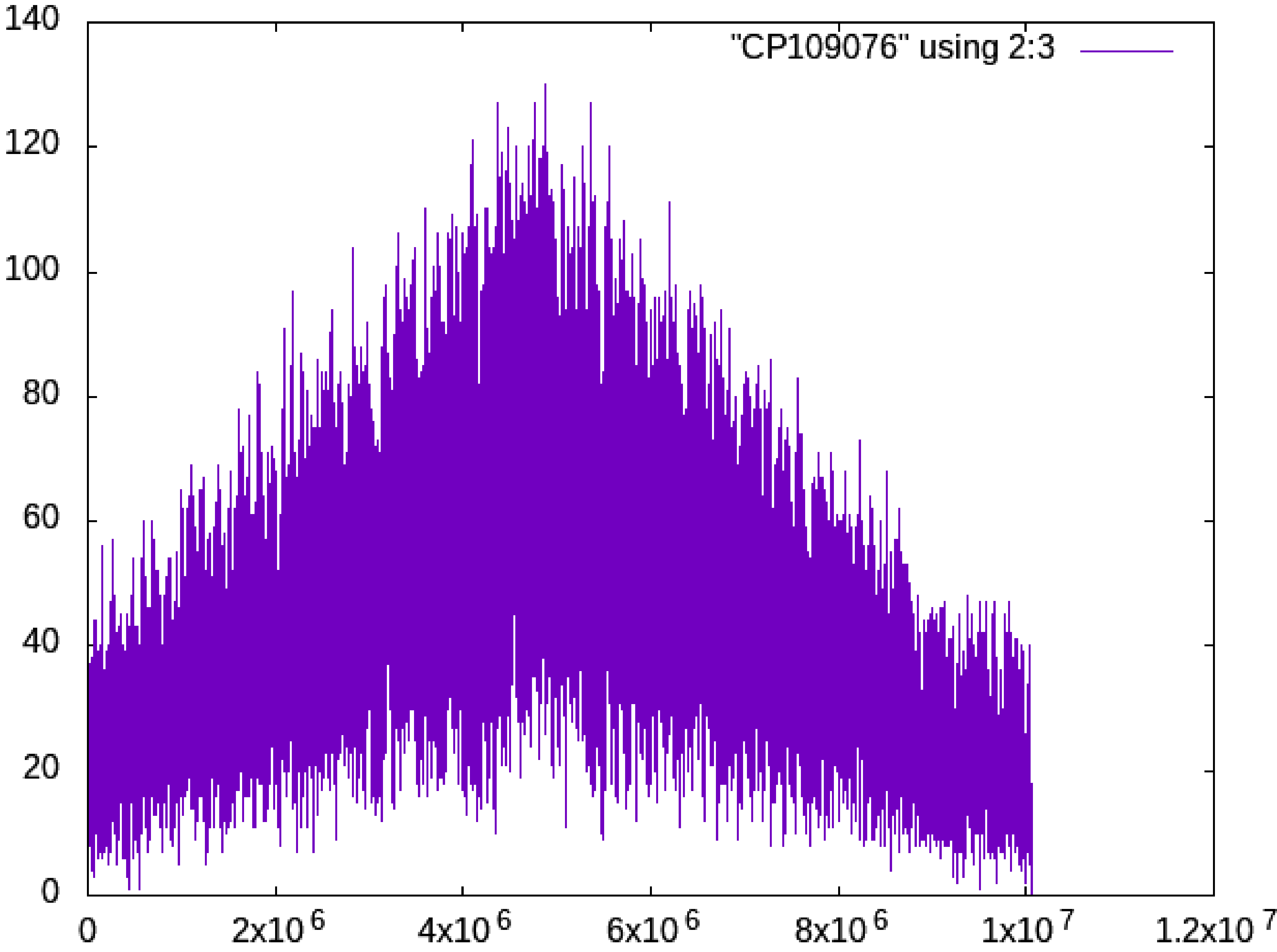

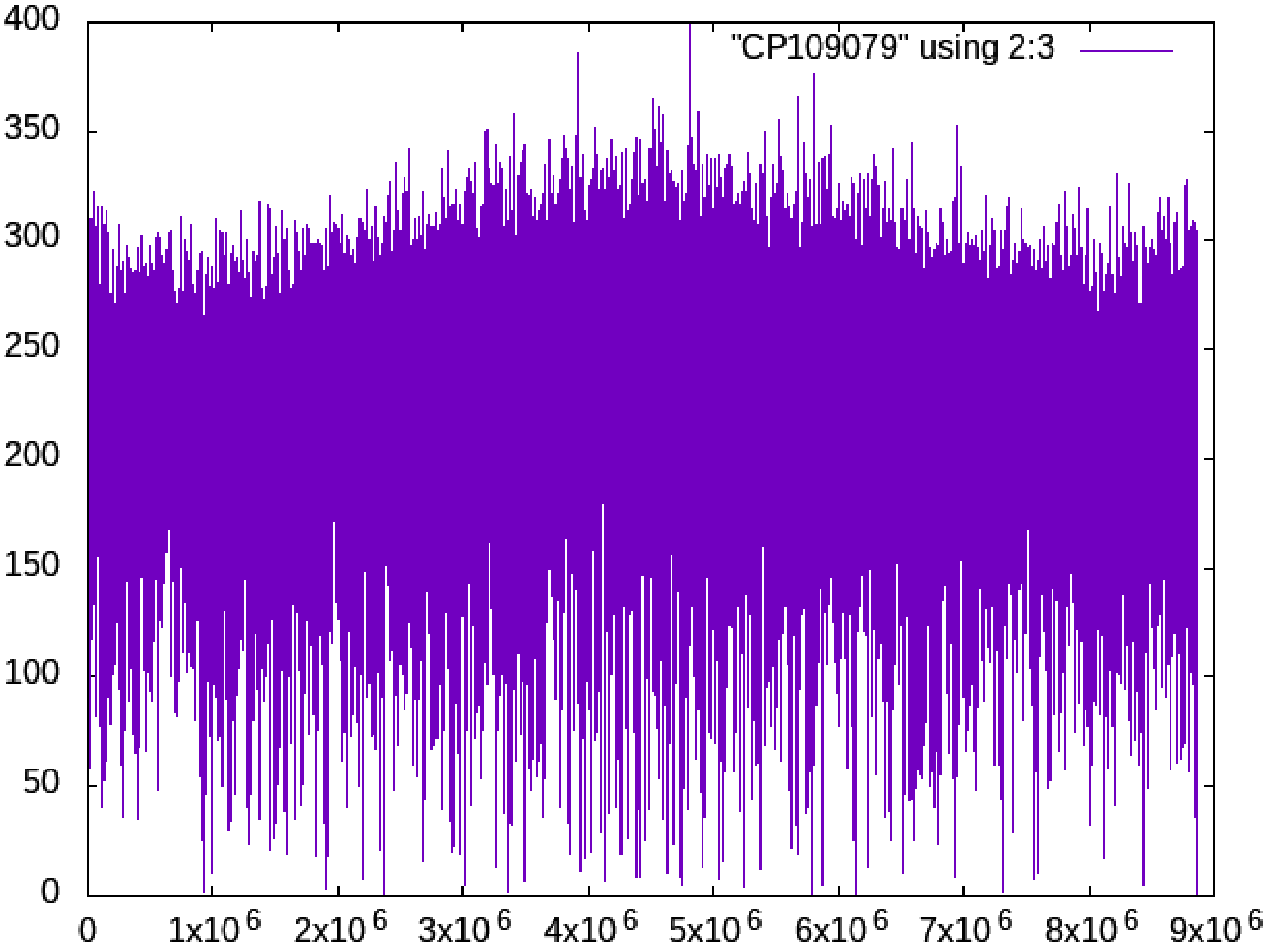

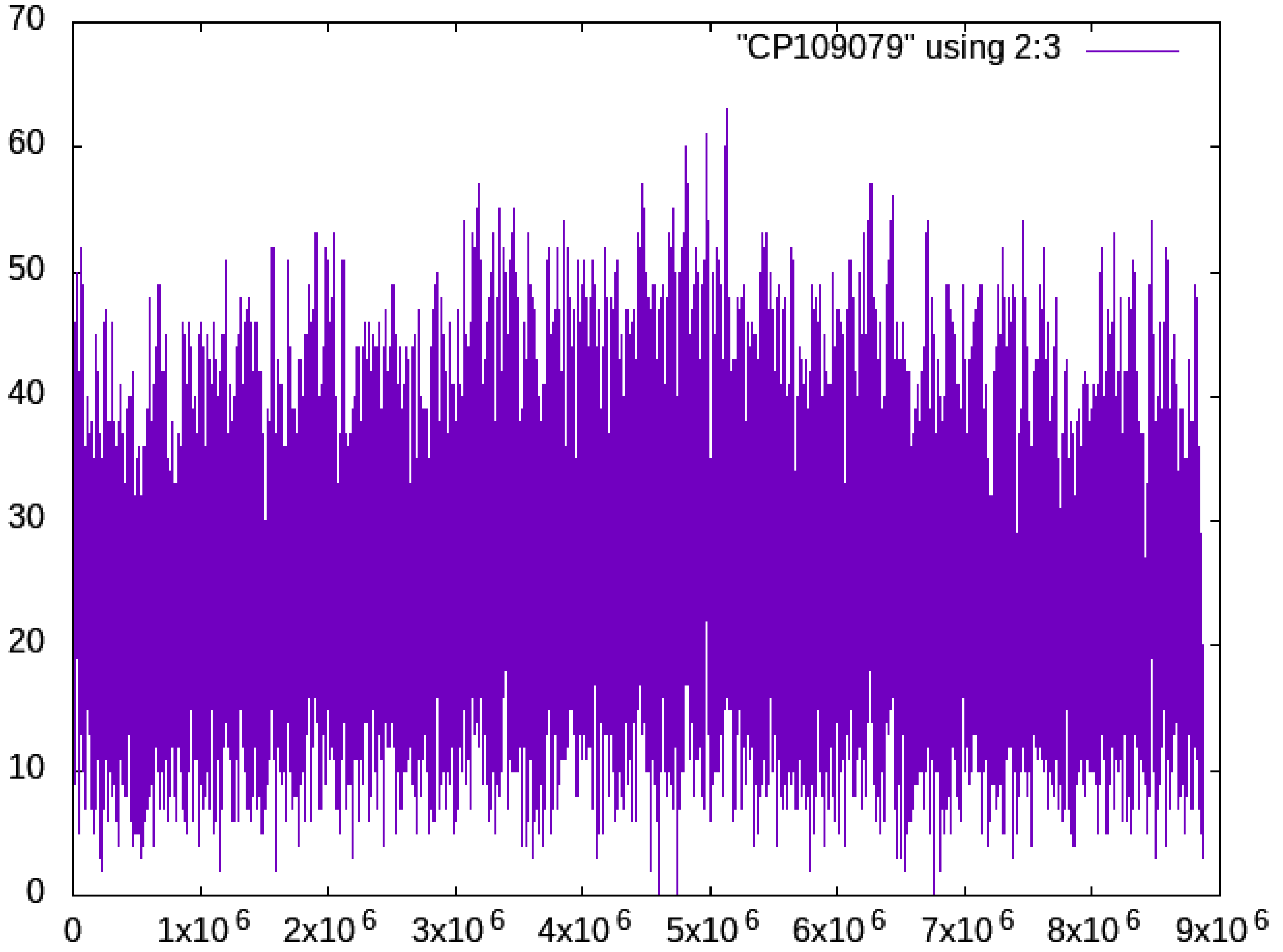

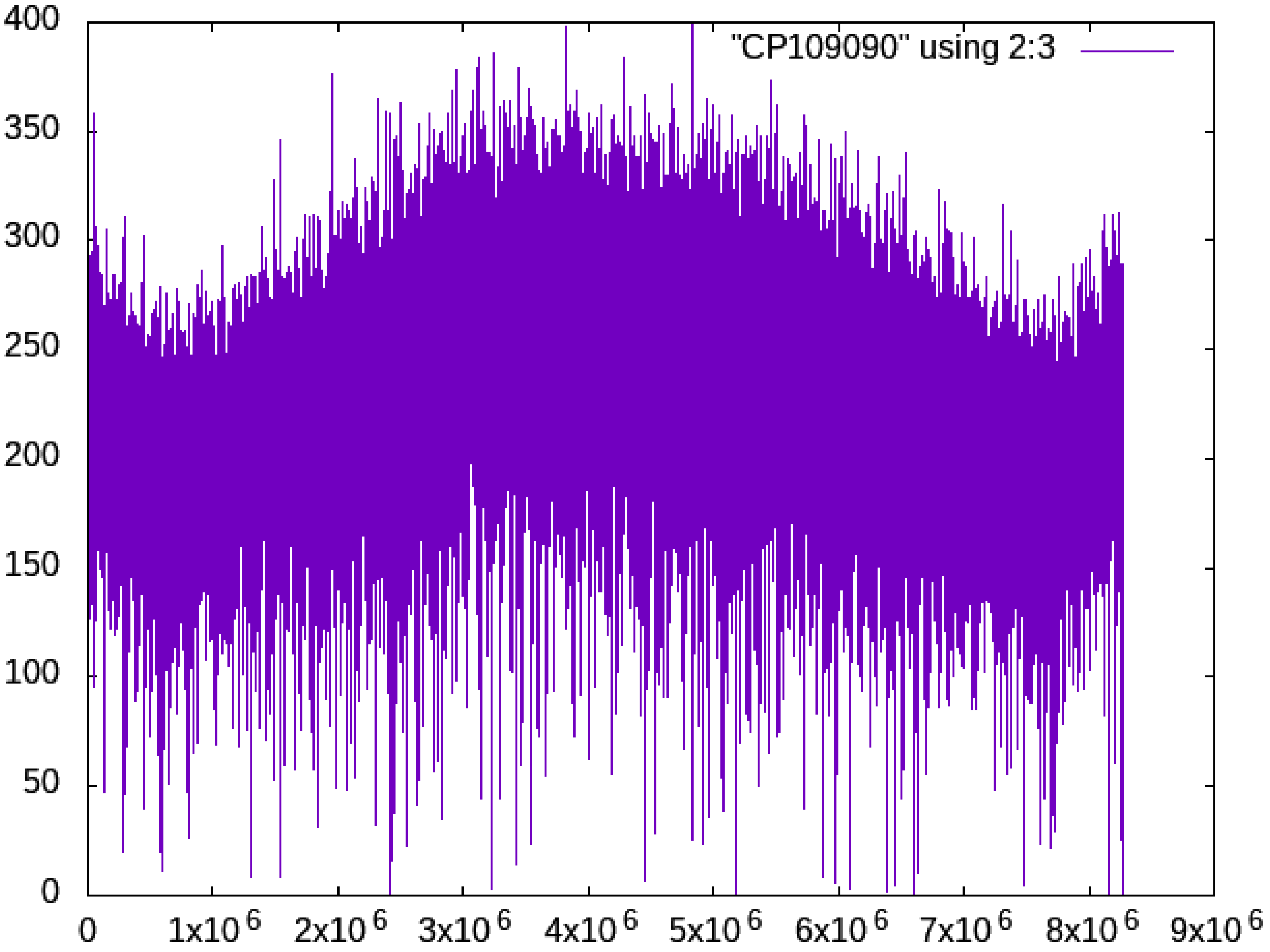

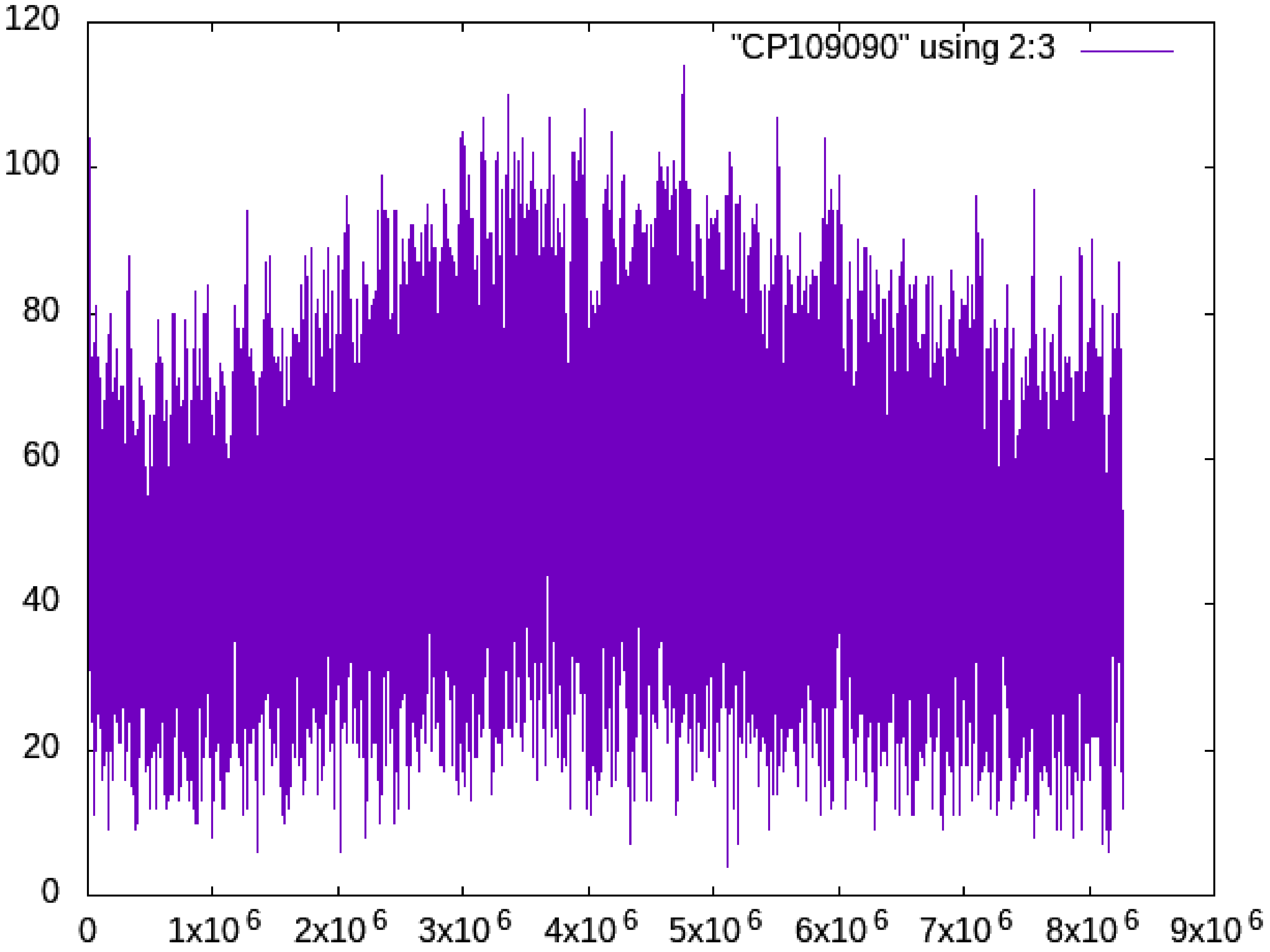

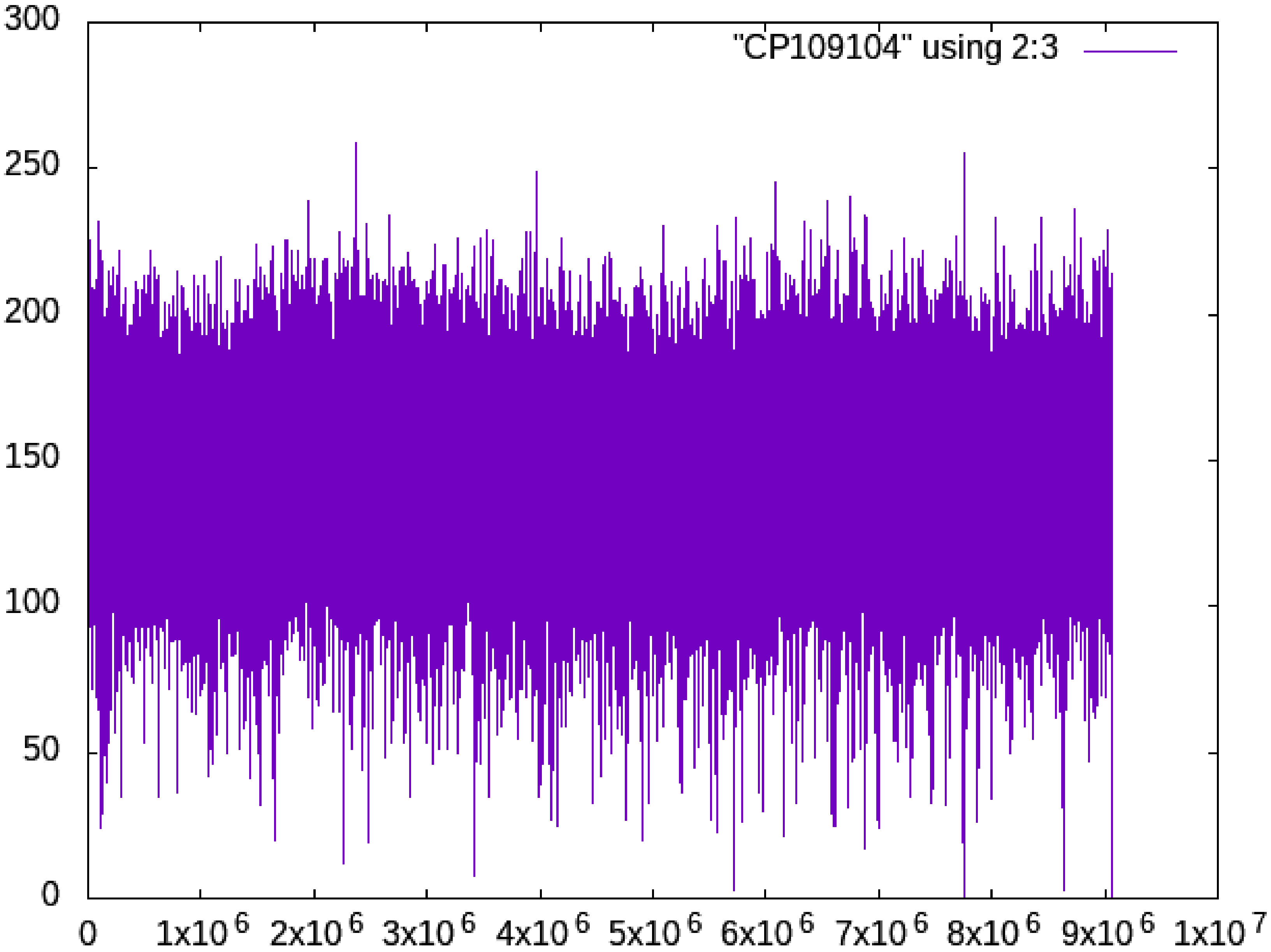

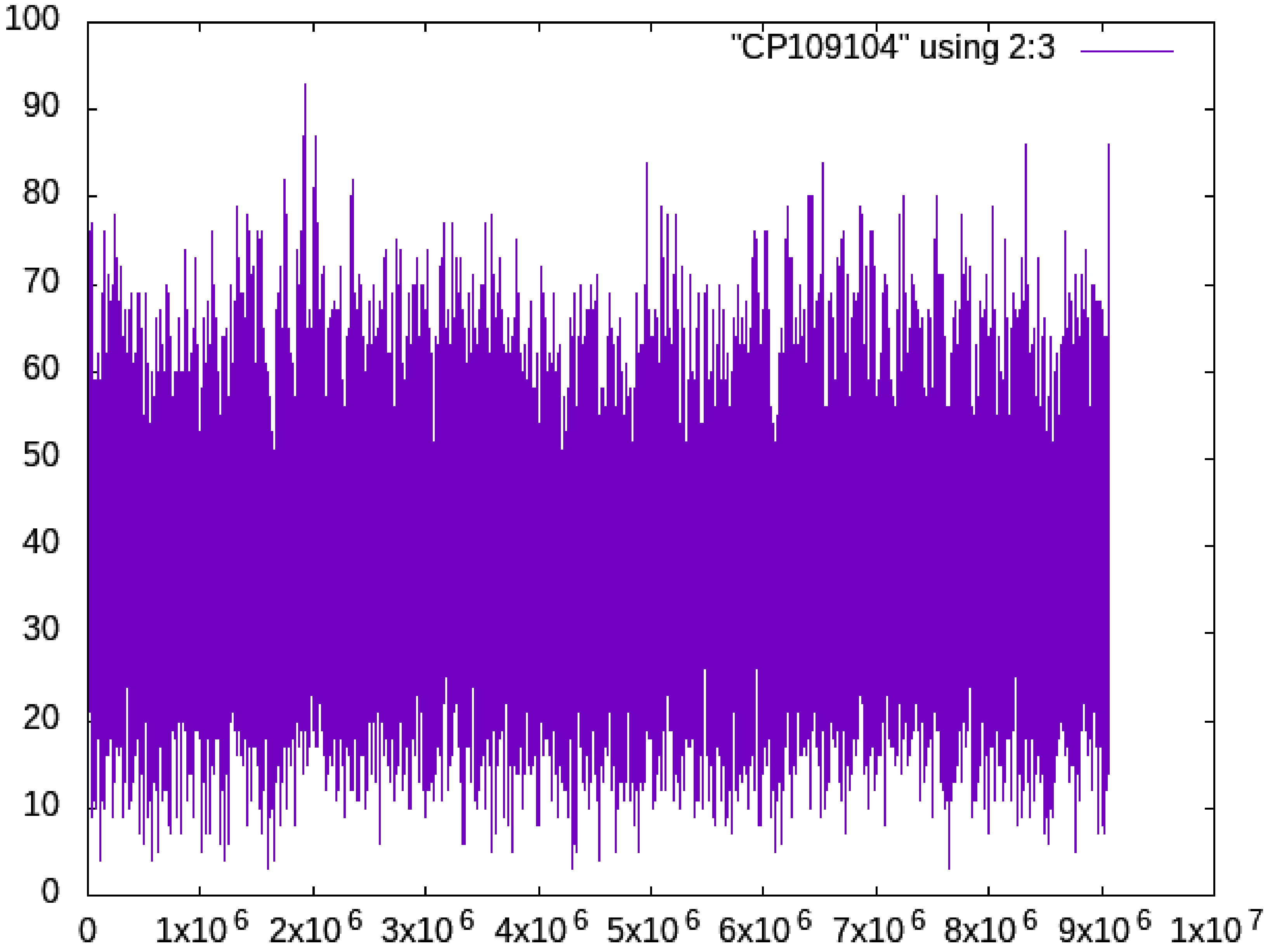

### Supplementary_material_S3_list_of_countries

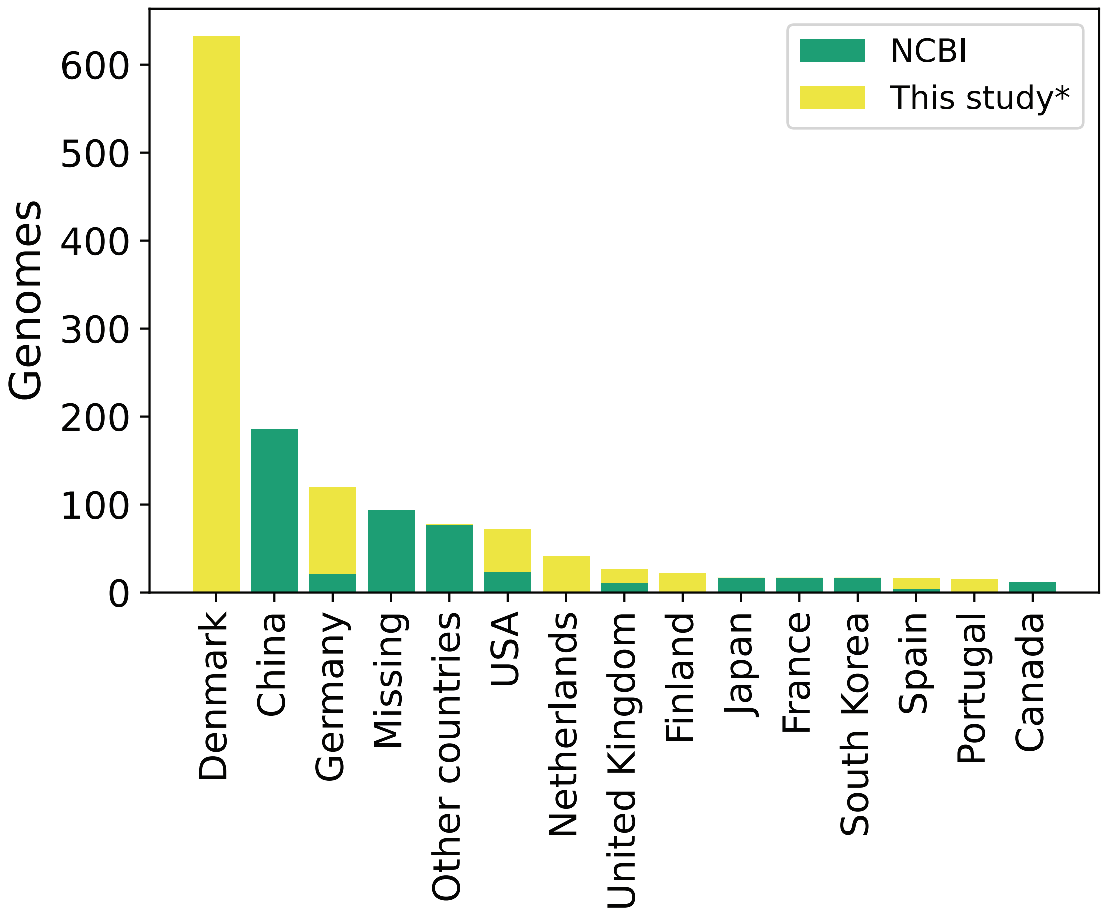

### Supplementary_material_S4_core_gene_placement_legend

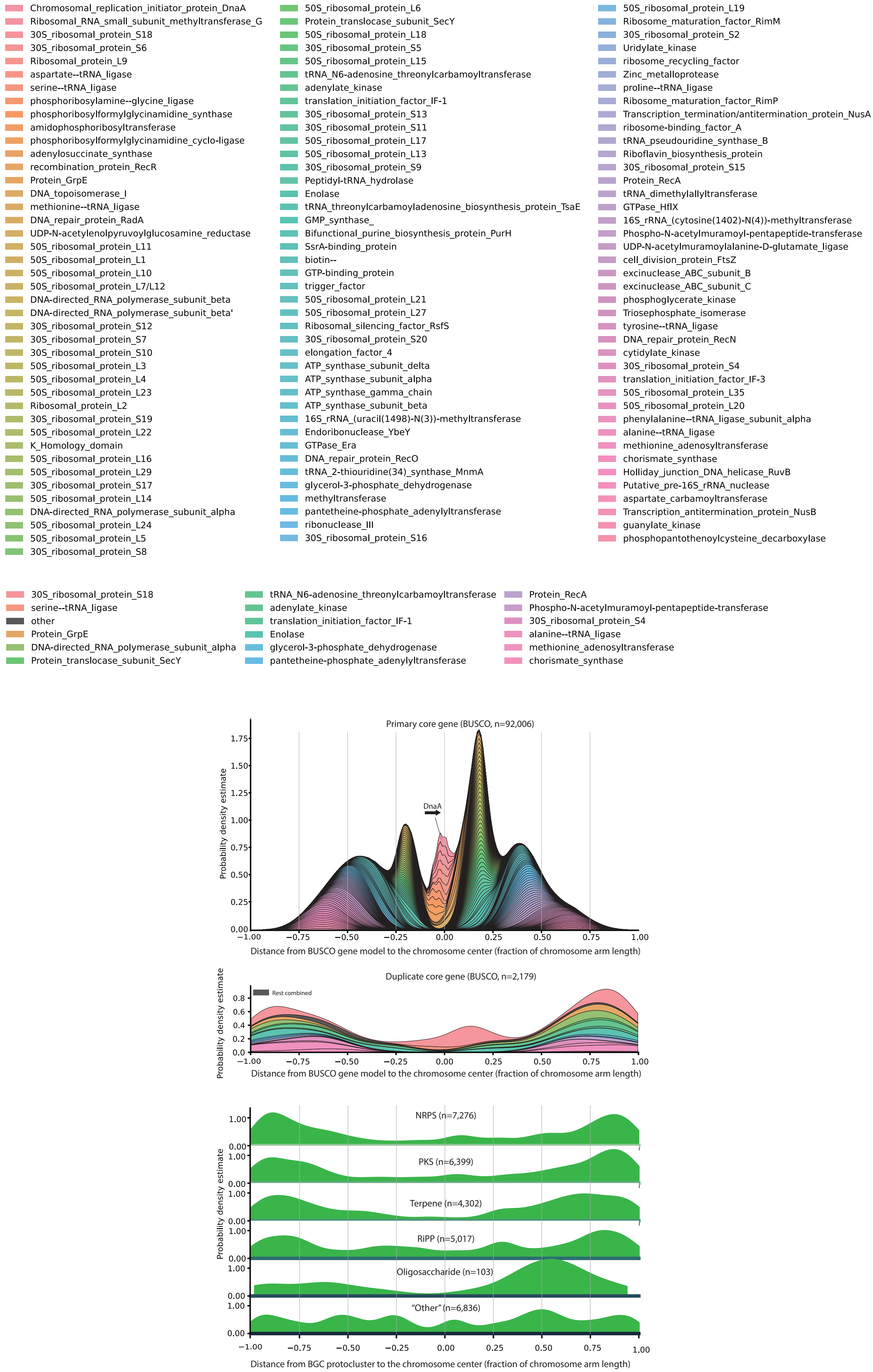

### Supplementary_material_S7_Antismash_cluster_type_plots_all_protocluster_types

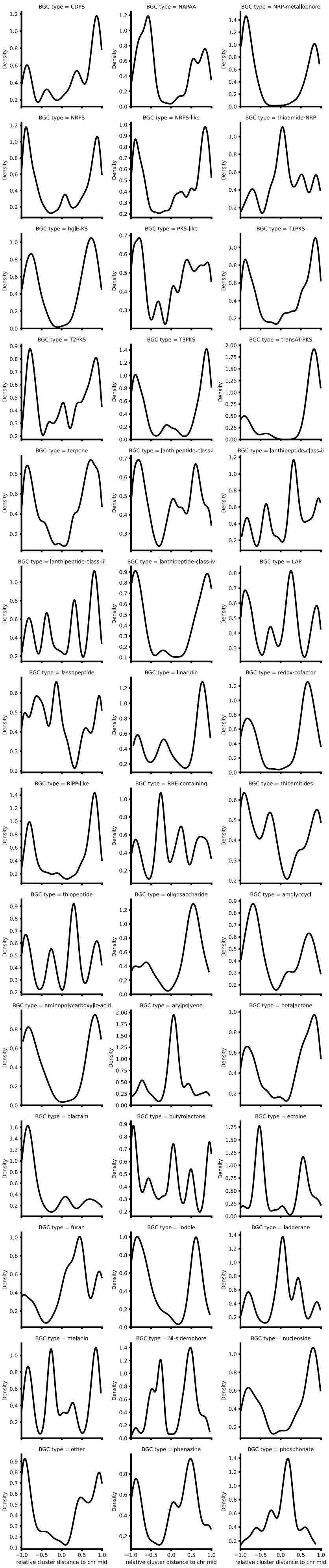

### Supplementary_material_S8_figure_clinkers_w_names

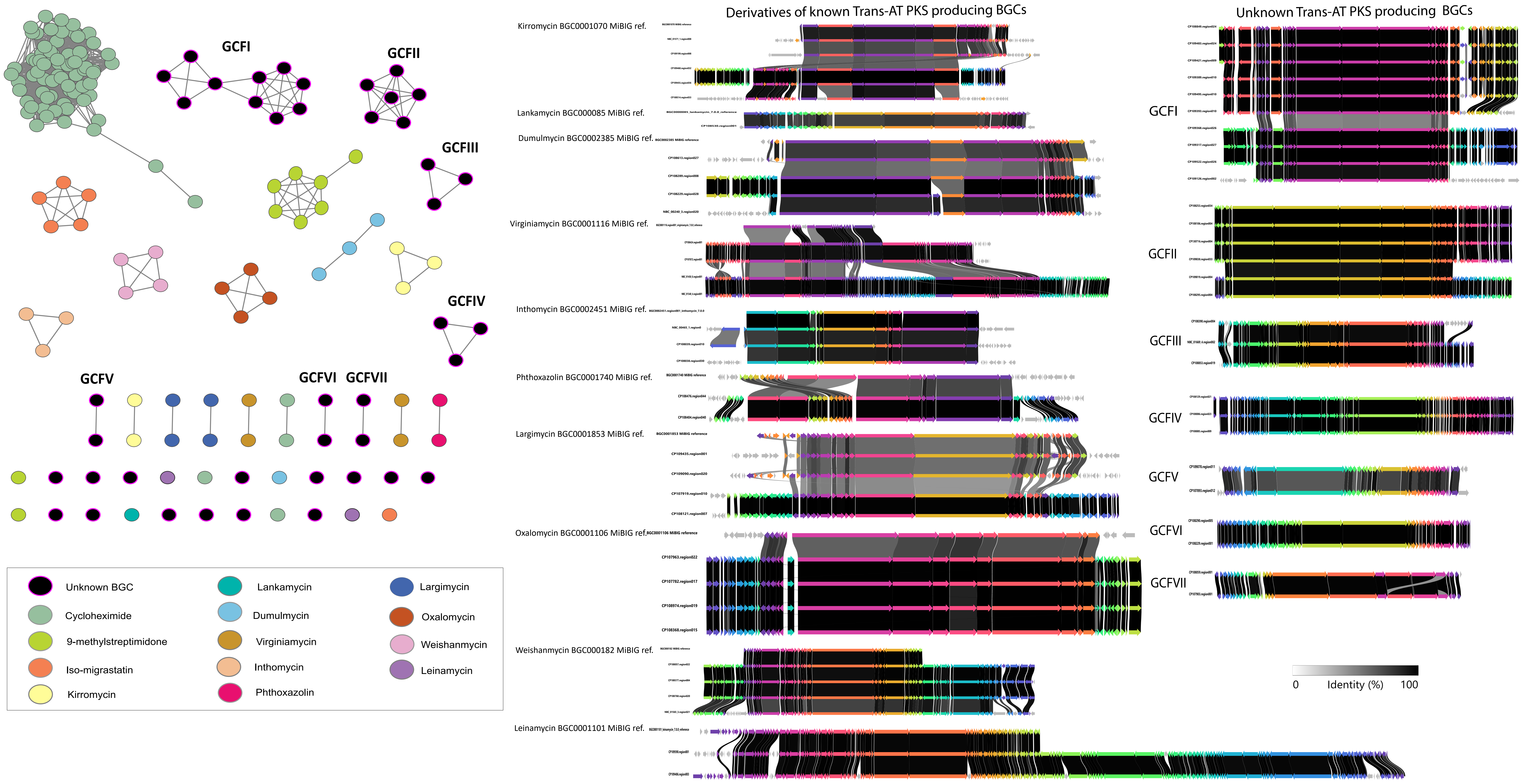
